# Photodisruption of the inner limiting membrane promotes retinal engraftment of stem-cell derived retinal ganglion cells

**DOI:** 10.1101/2025.04.14.648093

**Authors:** E. De Coster, K. De Clerck, C. De Clercq, W. Li, D. Punj, B. Vanmeerhaeghe, S. De Smedt, K. Braeckmans, H. Hadady, K. Remaut, T.V. Johnson, K. Peynshaert

**Affiliations:** Laboratory of General Biochemistry and Physical Pharmacy, Faculty of Pharmaceutical Sciences, Ghent University, Ghent, Belgium; Glaucoma Center of Excellence, Wilmer Eye Institute, Johns Hopkins University School of Medicine, Baltimore, MD, United States

**Keywords:** Glaucoma, ILM photodisruption, transplantation, stem cell therapy, collagenase

## Abstract

Glaucoma is the leading cause of irreversible blindness, driven by the progressive loss of retinal ganglion cells (RGCs). Stem cell-derived RGC transplantation could revolutionize glaucoma treatment, but the inner limiting membrane (ILM) remains a major obstacle by hindering cell migration into the retina. Interestingly, the ILM represents a double-edged sword for RGC engraftment: on the one hand, it greatly hinders cell migration, whereas on the other hand, its presence during retinal development is necessary for neuronal migration and retinal lamination. As an alternative to current invasive and harmful strategies to disrupt the ILM, we introduce ILM photodisruption, a minimally invasive biophotonic method that can manipulate the integrity of the ILM with unprecedented precision. In this study, we have finetuned the technology in bovine and human organotypic retinal explants to create templated ILM pores, creating entryways for donor RGCs to enter the retina while preserving most of the membrane to confer guidance cues for their engraftment. Applying this technology, we were able to promote donor RGC survival, enhance cell spreading and facilitate integration into the retina. Overall, our findings demonstrate that ILM photodisruption effectively addresses a key barrier in RGC replacement, paving the way for advancing retinal regeneration toward clinical application.

## 1. Introduction

Glaucoma is the leading cause of irreversible blindness worldwide, with an estimated global prevalence of 76 million individuals (1,2). As a consequence of our aging society, glaucoma prevalence is predicted to nearly double by 2040, which underscores the pressing need for more effective treatments (3,4). The primary hallmark of glaucoma and other optic neuropathies is the progressive and irreversible loss of retinal ganglion cells (RGCs). As the projection neurons of the retina, RGCs are solely responsible for transmitting visual information from the eye to the brain; a task that requires them to extend long axons through the optic nerves and tracts and synapse within subcortical targets of the visual system. Considering this essential function, it is not surprising that the loss of these neurons causes vision loss, and can eventually lead to total blindness (4,5).

Since elevated intraocular pressure (IOP) is the most important modifiable risk factor for glaucoma, current clinical treatments largely focus on delaying vision impairment by lowering the IOP through administration of drugs, laser treatments, or surgeries (4,6). Yet, many patients develop significant vision loss prior to diagnosis and lowering IOP does not always halt disease progression (4). Aware of these limitations, researchers are exploring alternative therapies such as neuroprotective strategies to prolong RGC survival and/or to provide support to the remaining neurons (4). Importantly, no existing glaucoma therapy can compensate for RGC death and hence reverse blindness. However, progress in the field of stem cell biology and neuroscience over the past 20 years has set the stage for exogenous replacement of lost neurons by transplantation of stem cell-derived RGCs to be feasible. This concept was substantiated by several pioneering studies which showed that intravitreally injected donor RGCs engrafted into the retina, acquired the correct morphology, and responded to light (5,7–9). While subsequent encouraging studies followed suit by substantiating the feasibility of RGC transplantation, they also revealed several challenges hampering its translation to human patients. Examples include the scaling up of RGC production and suboptimal functional connectivity of donor RGCs within the host retina circuitry (5,10–12).

Another dominant bottleneck hampering efficient RGC transplantation is poor migration of the donor RGCs from the vitreous into the retina due to the presence of the inner limiting membrane (ILM) which forms a physical border between the vitreous and the retina (Figure 1) (11,13,14). Its composition resembles that of a basement membrane, composed of collagen IV, laminin, glycoproteins, and heparan sulphate proteoglycans (15,16). Indeed, the majority of intravitreally injected cells accumulate at the ILM, resulting in negligible spontaneous cell migration into the retina (5,11,17). The ILM presents such a formidable barrier that some researchers have resorted to subretinal injection as an alternative method to deliver RGCs to the inner retina (18). Nevertheless, it is also important to note that basement membranes are more than “pretty fibrils” (19); they are now known to regulate many fundamental biological processes through membrane-cell interactions (20,21). The ILM is no exception. The ILM is essential during retinal histogenesis (22,23), where ILM-RGC interactions mediate coordinated patterning of the RGC layer as well as correct dendritic projection. Loss of ILM components, like laminin or fibronectin, results in defects in axon orientation and elicits the formation of RGC aggregates rather than the normal single cell layer (13,24). Interestingly, RGC transplantation shares similarities to retinal development, as donor RGC somata that arrive in the retina need instructive signals to orient themselves and their neurites effectively. Thus, ILM components may be essential to achieve functional integration within the visual neurocircuitry (11,13,25–27). This renders the role of the ILM toward RGC transplantation conflicting: on the one hand its components likely function as essential cues for RGC maturation and functional integration into the retina, yet on the other hand it serves as a physical barrier for cell entry, necessitating interventions for stem cells to effectively traverse it (27–30).

**Figure 1:**
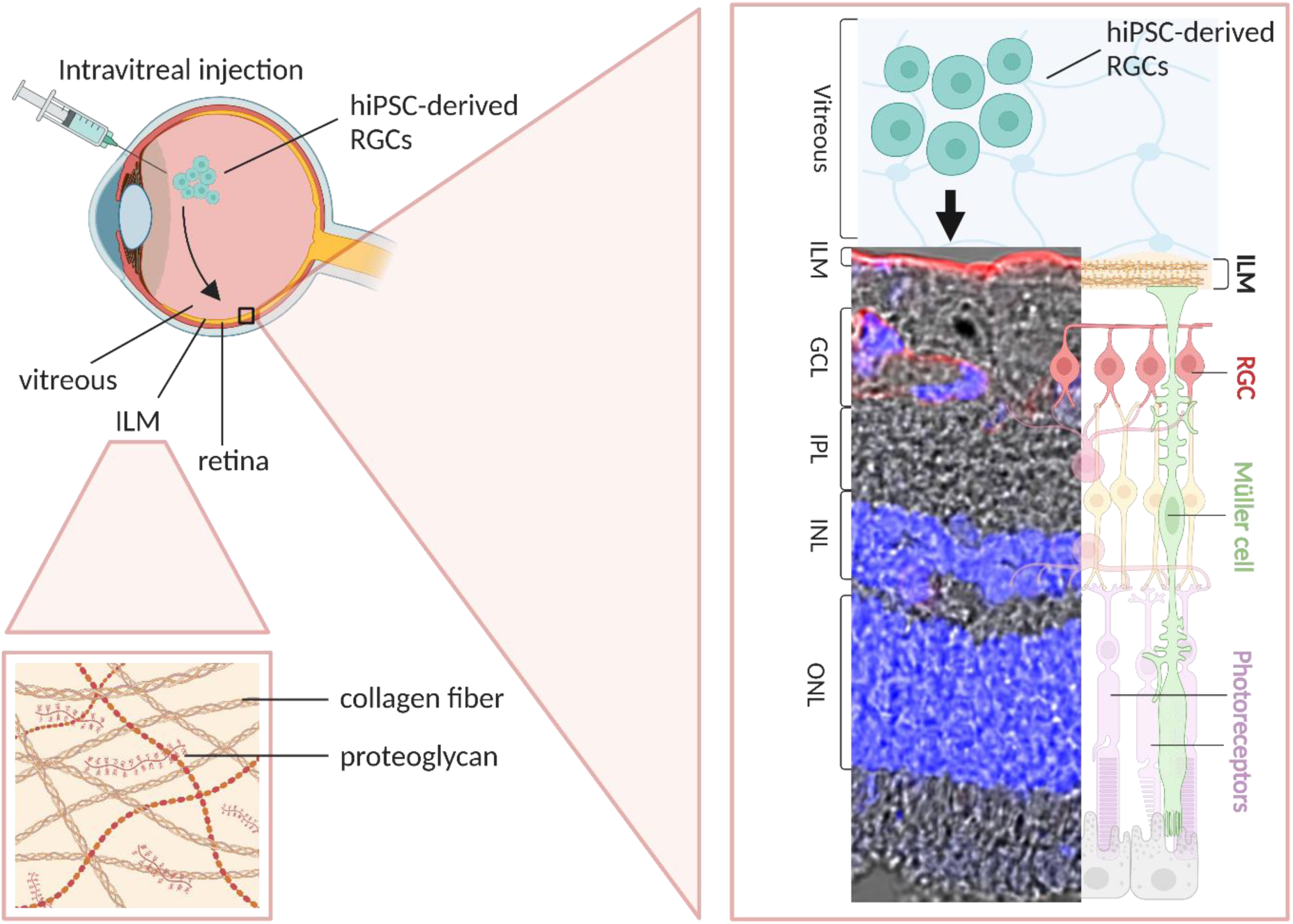
Schematic representation of the hampered migration donor RGC from the vitreous into the retina due to the presence of the ILM and confocal image (40x objective) of cryosection of untreated bovine retinal explant. The ILM and blood vessels were immunostained with laminin (red), all nuclei were stained with Hoechst (blue). Created with BioRender.com.

In context of this and the barrier role of the ILM for other therapeutic classes (16,31–33), several groups have explored strategies to disrupt the ILM or evade it altogether, including enzymatic digestion (30,34,35), ILM peeling (36) and mechanical disruption (17). Digestion of the ILM using enzymes like pronase has been shown to boost engraftment of stem cell-derived RGCs within the murine retina, by reducing clustering and increasing retinal dendritic ingrowth (30,37,38). However, the therapeutic window for enzymatic ILM degradation is narrow and retinal toxicity has been observed (13,30,34). In the context of retinal gene therapy, researchers aimed to enhance the retinal delivery of intravitreally injected adeno-associated vectors (AAVs) by surgically removing the ILM altogether (36,39). Although it is an established surgical technique used in human patients to treat vitreomacular traction and is highly effective in enhancing retinal transduction (17), ILM peeling is invasive and may harm the underlying structures, potentially limiting its application for RGC transplantation (40,41). Taken together, there has been no methodology to precisely modify the integrity of the ILM - until now.

As an alternative strategy we have developed a biophotonic approach to locally disrupt the ILM in a highly controlled manner based on ‘photoporation’, a concept we initially developed in our group for permeabilization of cell membranes to promote intracellular delivery of biologics (42–44). Our light-based approach relies on the creation of vapor nanobubbles (VNBs) induced by irradiating photosensitizers with high-intensity short laser pulses (45,46). First, the photosensitizer indocyanine green (ICG) is applied to the ILM surface of the retina, followed by application of extremely short laser pulses (<7 ns). This creates an ultrafast increase in temperature, resulting in evaporation of the surrounding water. Consequently, nanoscopic VNBs, are created which expand by consuming thermal energy. Finally, upon their collapse, high-pressure shock waves are released which can mechanically create pores in the ILM (Figure 2). For ILM photodisruption specifically, the FDA-approved organic dye indocyanine ICG is an ideal photosensitizer, as it is established in the ophthalmologic field as a dye to stain the ILM during surgical ILM peeling. Moreover, its favorable spectral properties operating in the near-infrared (NIR) range are beneficial in an *in vivo* setting (47).

**Figure 2:**
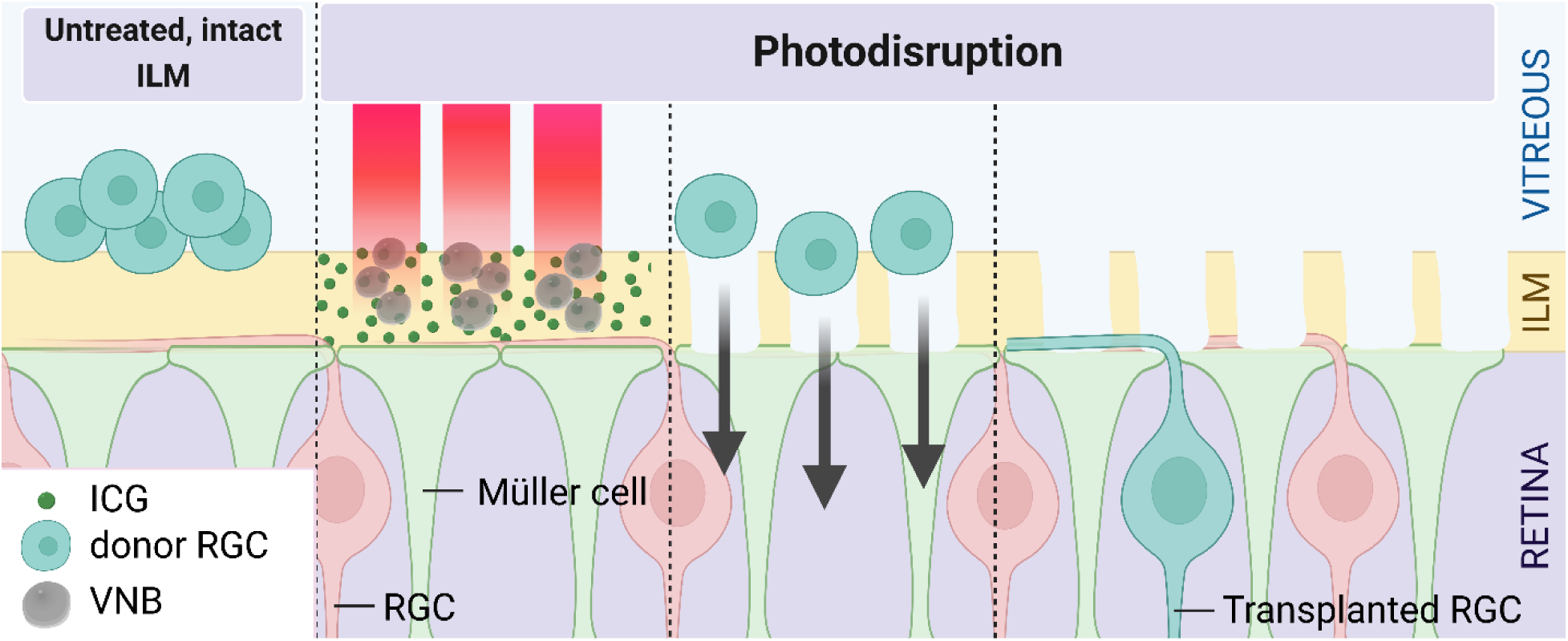
Schematic representation of ICG-mediated ILM photodisruption. Upon applying a nanosecond laser pulse, vapor nanobubbles (VNBs) emerge where ICG has accumulated at the ILM. The subsequent collapse of the VNBs induces local disruption of the ILM, creating pores and allowing the donor RGCs to enter the retina. This light-based technique features a lot of spatial control, enabling targeted disruptions of the ILM. Created with BioRender.com.

A powerful asset of (ICG-mediated) ILM photodisruption in comparison with the other reported strategies is tunability, as we have found that adjusting the ICG concentration and laser pulse energy allows us to control the extent of VNB formation and hence the degree of ILM disruption. Another important feature is its precise spatial control, enabling – for the first time – the targeting of specific retinal regions and even the creation of ILM breaks in a programmed pattern. We hypothesized that ILM photodisruption may be leveraged as a unique technique to controllably manipulate the ILM in such a way that it is overcome as a delivery barrier but leaves enough ILM intact to function as a guiding cue for newly transplanted RGCs. We therefore searched for a pattern that keeps most of the ILM intact, by varying the concentration of the photothermal agent ICG. To do so, we evaluated ILM integrity (i.e. pore diameter, pore area and percentage of intact ILM) following photodisruption in bovine retinal explants. In addition, we verified our observations on the thicker human ILM, thereby demonstrating species-specific differences in ILM response to photodisruption. Furthermore, we demonstrated the positive impact of our light-based approach on RGC transplantation in the bovine retina, evidenced by improved RGC survival, cell spreading, regularity, and neurite formation. We compared the effects of photodisruption to those of collagenase, an ILM-digesting enzyme currently considered the state-of-the-art method for ILM removal and used here as a positive control. Finally, we assessed the safety of the different approaches by examining retinal thinning, RNA-binding protein with multiple splicing (RBPMS) and gliotic markers.

## 2. Materials and methods

### Differentiation and purification of hiPSC-derived RGCs

RGCs were derived from human induced pluripotent stem cells (hiPSCs) carrying genes for tdTomato and the murine cell-surface protein CD90.2/THY1.2 driven by the endogenous POU4F2 (BRN3B) promoter, as described by Sluch, et al. (48–51). Briefly, the episomal derived EP1 hiPSC line (50,52) was maintained by clonal propagation in mTeSR™ 1 medium (Stem Cell Technologies™) on growth factor-reduced Matrigel-coated (Corning™) 6-well plates in an incubator at 37°C in 5 % CO_2_. At day 0, hiPSCs were replated on new Matrigel-coated plates (Corning™) in mTeSR™ Plus medium (StemCell Technologies™) followed by replacement of this medium on day 1 by supplemented Neurobasal A medium. Next, to initiate and promote differentiation, a set of small molecules was added to the cells in a well-established time schedule for a period of up to 40 days. The working concentrations of small molecules were: Forskolin (25 µM, Millipore-Sigma), Dorsomorphin (1 µM, Stem Cell Technologies™), Inducer of definitive endoderm 2 (2.5 µM, R&D Systems), DAPT (10µM, Millipore-Sigma), and Nicotinamide (10 mM, Millipore-Sigma). Following differentiation, RGCs were isolated using magnetic activated cell sorting (MACS) by targeting the surface antigen Thy 1.2 (30,48). The same batch of hiPSC-derived RGCs was used for all experiments described in this paper.

### Bovine retinal explant dissection

The protocol of bovine explant dissection was performed as described previously (45), minor alternations are listed below. Briefly, fresh bovine eyes were obtained from a local slaughterhouse and transported in cold CO_2_ independent medium (Gibco™). After discarding all extra-ocular tissue and disinfecting the eyes in 20% ethanol, the anterior segment and the vitreous were removed. While the posterior segment was submerged in cold CO_2_ independent medium, the eye cup was flattened by making 3 relaxing incisions. To isolate 11 mm explants for laser treatment, a corneal trephine blade of 11 mm in diameter (Beaver®) was used, whereas a 5 mm trephine (Electron Microscopy Sciences™) was utilized to obtain 5 mm explants for cryosectioning or flatmounts. The optimal pore diameter, pore area and percentage of intact ILM were assessed across 13 different bovine retinal explants. The transplantation experiment with 20.000 RGCs was conducted on 15 retinal explants, while the experiment with 33.000 RGCs was performed on 21 retinal explants.

### Human retinal explant dissection

Human retinal explants were prepared according to our previously published protocol (45). Postmortem human eyes were obtained from Biobank Antwerpen, (Antwerp, Belgium; ID: BE 71030031000) and Ghent University Hospital. Protocols were approved by the Ethical Committee of Ghent University Hospital (dossier number: B670201837281; EC/2018/1071). Eyes were kept in CO_2_ independent medium (Gibco™) at 4°C until dissection. First, the anterior segment was removed by cutting through the sclera and the vitreous. Using forceps, the eyecup was placed upside down with the optic nerve upwards. While holding the optic nerve region tight with forceps, the optic nerve area was then excised from the outside. Next, the retina with vitreous attached - was removed from the eyecup and flattened using a soft paintbrush. Several 5 mm diameter retinal explants were then punched out using a corneal trephine blade of 5 mm in diameter (Electron Microscopy Sciences™). In particular, 5 mm explants were derived from the near peripheral region where the ILM is at its thickest, thus avoiding the thin ILM in the region of the fovea centralis (53). Subsequently, the vitreous was removed by sticking a dry piece of filter paper to the photoreceptor side of the retina after which the vitreous could be gently removed using forceps. Subsequently, the 5 mm diameter explant was detached from the filter paper by placing it in cold CO_2_ independent medium. The human retinal explants investigated in the photodisruption experiments were from 5 different donors aged 65-78y (i.e. male 65y, female 72y, female 74y, male 76 y, and male 78 y), whereas the human retinal explants for the collagenase experiments were obtained from 3 different donors (male 49y, male 70y and male 75y). Both eyes of each human donor were examined in this study.

### Laser treatment of bovine and human explants

Indocyanine Green (Merck™) was dissolved in distilled water at concentrations ranging from 0.10 mg/ml to 1.00 mg/ml and protected from light until use. A fresh solution was prepared for each experiment. Based on pore size optimization experiments with varying ICG concentrations, we determined that an ICG concentration of 0.25 mg/ml was optimal and used this concentration for all hiPSC-RGC transplantation experiments. Retinal explants with a diameter of 11 mm, were transferred into a 35mm glass bottom dish (Nunc™) with the ILM surface upwards. Next, the dish was placed in the optical set-up and 20 µl of ICG solution was applied to the top of the explant. Laser treatment was performed immediately after ICG application. A custom-made optical setup, built around an inverted TE2000 epi-fluorescence microscope (Nikon), was used to generate and detect VNBs. In this set-up, a 7 ns pulsed laser with a laser beam size of 92 μm was tuned to a wavelength of 800 nm (Opolette HE 355 LD, OPOTEK Inc.). The energy of the laser pulses was monitored with an energy meter (J-25MB-HE&LE, Energy Max-USB/RS sensors, Coherent). In all experiments, the same laser fluence of 1.65 J/cm^2^ was applied. An automatic Prior Proscan III stage (Prior scientific Ltd.) was used to scan the sample through the distinct photodisruption pattern with a scanning speed of 2500 µm/s and an electronic pulse generator (BNC575, Berkeley Nucleonics Corporation) was applied to fire single laser pulses. The total scanning time of one 11 mm retinal explant was approximately 15 minutes. Immediately, after laser treatment, two 5 mm diameter explants were punched out of each 11 mm explant. Then, explants were either fixed immediately or cultured for 8 days with ILM side upwards.

### Collagenase treatment of bovine and human explants

Collagenase type I (Sigma Aldrich - product number C0130 - batch number SLCN7779) was diluted in phosphate-buffered saline (PBS) (without calcium and magnesium) to reach the desired activities (varying from 50 U/ml to 200 U/ml). The activity reported by the manufacturer was checked using an EnzChek™ Gelatinase/Collagenase Assay Kit. The same batch of collagenase was used for all experiments. For the RGC transplantation experiments, 5 µl of 50 U/ml collagenase was applied to the ILM surface of each 5 mm explant. After 24 hours of incubation at 37°C and 5% CO_2_, the collagenase solution was washed away with an excess of PBS. Afterwards, explants were fixed immediately or placed into culture for 7 more days.

### Culturing bovine and human explants

The 5 mm explants were cultured with the ILM side upwards in culture inserts (Millipore) in 6-well plates with retinal culture medium composed of Neurobasal-A, N2-supplement, B27-supplement, glutaMAX supplement, and penicillin-streptomycin (54). Half of the culture medium (i.e. 1.5 ml) was exchanged every other day. Explants were cultured at 37°C and 5% CO_2._

### Retinal ganglion cell transplantation in bovine explants

One day after photodisruption or collagenase treatment, hiPSC-derived RGCs were thawed and mixed with retinal culture medium to obtain a single cell suspension of 20.000 or 33.000 RGCs per 3 µl. This volume of 3 µl cell suspension was pipetted onto the ILM surface of each 5 mm diameter bovine explant and was cocultured for 7 days, as illustrated in Figure 3. In a 96-well plate coated with laminin and poly-L-ornithine (PLO) (54), 8000 RGCs (from the same vial as the transplantation) were cultured for 7 days to compare RGC survival and clustering.

**Figure 3:**
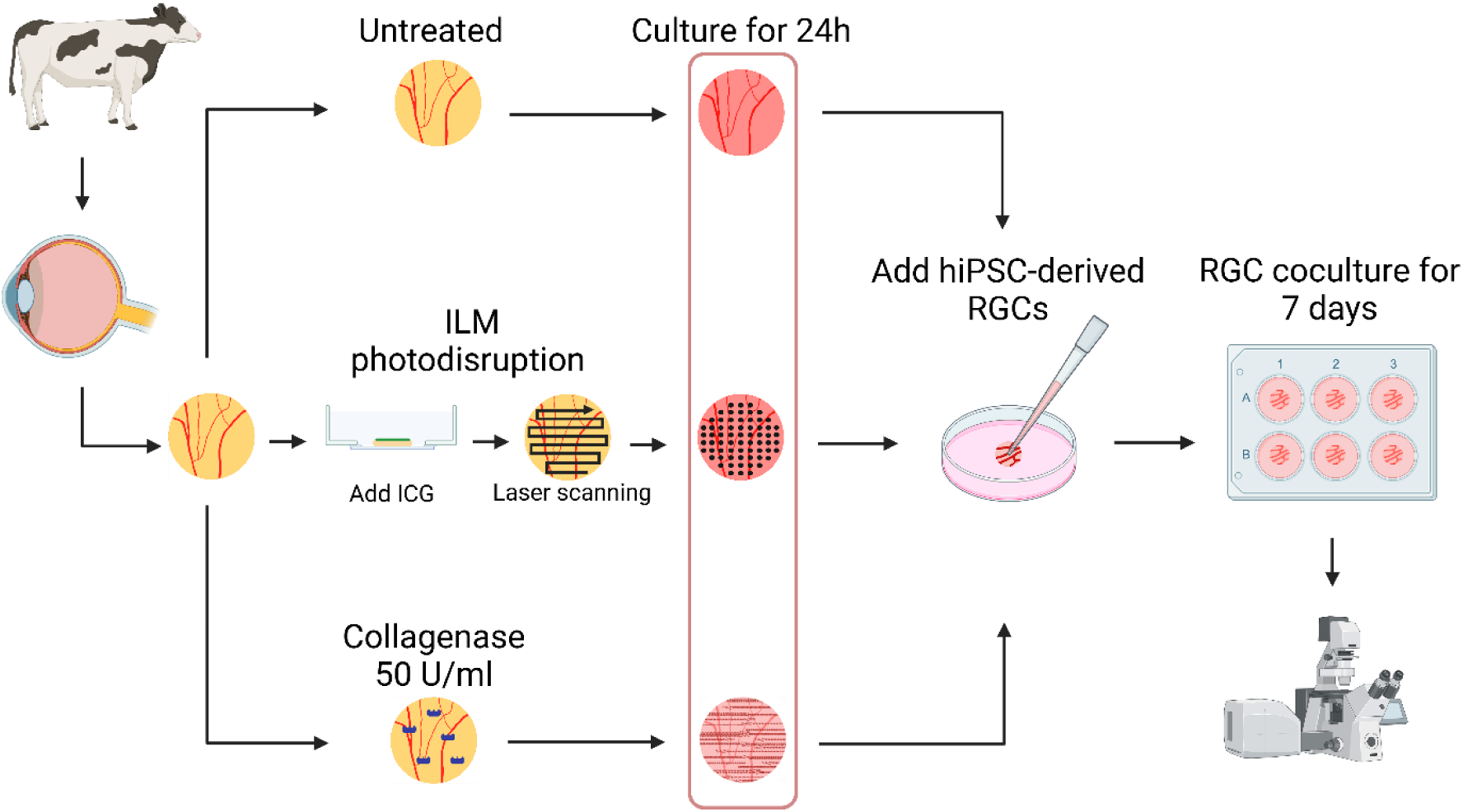
Schematic representation of the experimental design. The explants were treated (i.e. photodisruption or collagenase treatment) immediately following isolation. In pink, the integrity of the ILM is visualized. On the ILM side of each pre-treated bovine retinal flatmount, 20.000 or 33.000 hiPSC-derived RGCs were transplanted. Donor RGCs were cocultured for 7 days at 37°C followed by fixation, immunostaining and confocal imaging. Created with BioRender.com.

### Immunohistochemistry on bovine and human flatmounts

After treatment and/or (co-)culture, immunohistochemistry was performed on the retinal flatmounts according to previously described methods (30,54,55). Briefly, retinal flatmounts were fixed with 4% paraformaldehyde (PFA) at 4°C for 2h followed by a washing step with PBS. Next, the explants were blocked and permeabilized at room temperature for 1h using blocking buffer (0.3% triton, 10% goat serum in PBS). Subsequently, the retinas were incubated with primary antibodies diluted in blocking buffer for 5 nights at 4°C while shaking. Following washing with PBS, the flatmounts were incubated with the corresponding goat secondary antibodies and Hoechst overnight at 4°C on a shaker (Table 1). After final washing with PBS, the flatmounts were mounted on glass slides using 1:1 PBS-Glycerol as mounting medium. Flatmount imaging was performed with a Nikon A1R Confocal microscope applying a 10x air and a 40x air objective (plan apo λ 40X, NA 0.9, WD 250 µm). For the calculation of donor neuron survival, all donor RGCs within each explant were imaged and quantified, with a tiled Z-stack of the 5 mm flatmount, rather than sampling and extrapolating the survival rate. All image tiles were stitched to form one single image per flatmount. For the detailed pictures, random regions in the center of the explant were imaged with a 60x water objective (SR plan apo IR 60X WI, NA 1.3, WD 180 µm).

**Table 1:** Antibodies for flatmount staining used in this study indicating antigen target, host species, fluorophore, dilution, manufacturer, and identifier.

| Antigen target | Host species | Fluorophore | Dilution | Manufacturer | Identifier |
| --- | --- | --- | --- | --- | --- |
| GS | Mouse | / | 1:200 | Abcam | Cat#ab64613 |
| Human Nuclear antigen | Mouse | / | 1:300 | Sigma | Cat#MAB1281 |
| Laminin | Rabbit | / | 1:50 | Sigma | Cat#L9393 |
| RFP, cross-reactive to tdTomato | Rabbit | / | 1:300 | Rockland | Cat#600-401-379 |
| RBPMS | Mouse | / | 1:200 | Invitrogen | Cat#OTI3B7 |
| Mouse | Goat | Alexa Fluor 488 | 1:1000 | Invitrogen | Cat#A28175 |
| Rabbit | Goat | Alexa Fluor 568 | 1:1000 | Invitrogen | Cat#A-11036 |
| Rabbit | Goat | Alexa Fluor 647 | 1:500 | Invitrogen | Cat#A-21245 |
| Hoechst | / | Hoechst 33342 | 1:500 | Invitrogen | Cat#H3570 |

### Cryosection preparation and staining of bovine explants

After treatment and/or (co-)culture, retinal explants were fixed with 4% PFA at 4°C for 2h followed by cryopreservation, snap freezing, sectioning and staining (33,45). Then, the explants were submerged in 30% sucrose (overnight, 4°C); followed by a 1:1 solution of O.C.T (Tissue-Tek®)/30% sucrose (3 h, 4°C). Subsequently, the explants were embedded in pure O.C.T. (3h, room temperature) and snap frozen in isopentane cooled with dry ice. The frozen tissue blocks with bovine retinas were cut at 25 μm thickness using a Leica Cryostat and mounted on SuperFrost® Plus slides (Thermo Fisher Scientific). Cryosections were washed with PBS for 10 min followed by permeabilization for 5 min with 0.1% Triton™ X-100 (Sigma). After another washing step with PBS, sections were blocked in 5% goat serum (Thermo Fischer Scientific) for 1h. Next, sections were incubated overnight in primary antibody solution at 4°C (Table 2). Subsequently, the slides were washed with PBS and counterstained with the species-specific secondary goat antibodies and Hoechst for 1h at room temperature. Finally, the sections were washed one more time with PBS before cover slipping over mounting medium (1:1, PBS-Glycerol). Cryosection imaging was performed with a Nikon A1R Confocal microscope applying a 40x air objective (plan apo λ 40X, NA 0.9, WD 250 µm).

**Table 2:** Antibodies for cryosection staining used in this study indicating antigen target, host species, fluorophore, dilution, manufacturer, and identifier.

| Antigen target | Host species | Fluorophore | Dilution | Manufacturer | Identifier |
| --- | --- | --- | --- | --- | --- |
| GFAP | Rat | / | 1:1000 | Invitrogen | Cat#13-0300 |
| RFP, cross-reactive to tdTomato | Rabbit | / | 1:500 | Rockland | Cat#600-401-379 |
| Rat | Goat | Alexa Fluor 488 | 1:500 | Invitrogen | Cat# A-11006 |
| Mouse | Goat | Alexa Fluor 488 | 1:500 | Invitrogen | Cat#A28175 |
| Rabbit | Goat | Alexa Fluor 488 | 1:500 | Invitrogen | Cat#A-11034 |
| Rabbit | Goat | Alexa Fluor 568 | 1:500 | Invitrogen | Cat#A-11036 |
| Rabbit | Goat | Alexa Fluor 647 | 1:500 | Invitrogen | Cat#A-21245 |
| Hoechst | / | Hoechst 33342 | 1:500 | Invitrogen | Cat#H3570 |

### Image analysis and statistics

Pore diameter and pore area were determined by manually delineating the pores in Fiji using 10x and 40x objective images of bovine and human flatmounts. Subsequently, the percentage of intact ILM was calculated by subtracting the total area of all pores, representing the non-intact fraction, from the total area of each image. A total of 22 images were analyzed to determine the pore diameter of the bovine explants, whereas 16 human flatmount images from 5 different donors (aged 65-78y) were examined. Furthermore, Fiji was used to manually count the number of integrated donor hiPSC-RGCs in 3D and analyze their topographical localization within the explants; from which the percentage of cell survival and density were calculated. Only if positive for both tdTomato (red) and Human Nuclei (green), a donor RGC was counted.

Within each explant, all donor RGCs in the tiled Z-stack of the 5 mm flatmount were quantified. Density heat maps were generated using a customized protocol written in MATLAB (version R2023b). From each transplantation experiment, at least 1 explant per group was cryopreserved and imaged as cryosections.

Using the coordinates of the transplanted RGCs, the nearest neighbor distance (NND) was determined in Fiji (30,56). The NND measures the distance between a reference cell and its closest neighboring cell. As a result, each transplanted RGC has a single NND value; when combining all the NND values of an explant, the nearest neighbor index (NNI) can be calculated using Equation 1. This spatial metric provides insights into the relative spacing among neurons, which is indicative of cell clustering. In addition, the regularity index (RI) was calculated following Equation 2 (30,54,57,58). The RI is a measure of spatial uniformity, with higher RI values indicating less random and more uniform organization.

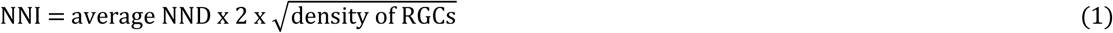

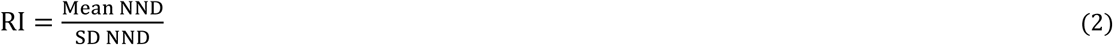

The number of neurites per cell in each cell layer were quantified manually in high magnification (60x objective) Z stacks using Fiji. Different cell layers were distinguished based on differences in intensity of the Hoechst signal of the bovine endogenous retinal cells using an orthogonal view or 3D viewer. The number of neurites per mm^3^ was determined by dividing the total number of neurites in each cell layer (IPL, INL and ONL) by the volume from the ILM to this specific cell layer. To obtain the number of neurites per mm^3^ per RGC, this value was further divided by the total number of RGCs in the Z-stack image. In the transplantation experiment with 20.000 RGCS, 1 detailed Z-stack was captured of each explant, whereas in the experiment with 33.000 RGCs, 2 Z-stacks were obtained per explant. Neurites in high magnification (60x objective) Z stacks were semi-automatically traced by a second independent, masked investigator using the Imaris - Autopath Filament tracer (v10.1, Oxford Instruments). Autodiameter feature was enabled to dynamically adjust for changes in neurite thickness along their length. This analysis measured the total area of neurites, neurite area per cell layer, total neurite length, neurite length per cell layer, and the branch level. Subsequently, retinal thinning was measured manually in Fiji by examining the difference in Z position between the different layers (ILM to IPL, ILM to INL, and ILM to ONL) on high magnification (60x objective) detailed Z stacks.

To quantify the number of endogenous RGCs, these cells were immunohistochemically labeled with antibodies against RNA-binding protein with multiple splicing (RBPMS). Next, tiled Z stacks of the whole flatmounts were manually counted using Fiji. Within each explant, all endogenous RGCs in the tiled Z-stack of the 5 mm flatmount were quantified. A total of 8 explants were analyzed (i.e. 2 explants from 2 independent bovine eyes per condition). The NND, NNI, and RI values of the untreated 0h flatmounts (i.e. flatmounts fixated immediately after dissection) were calculated based on the coordinates of the endogenous RGCs, as previously described for the transplanted RGCs.

Pooled data of both transplantation experiments were reported as mean ± SD unless otherwise stated. Group means were compared using one-way ANOVA with Tukey’s Post hoc test, unless otherwise stated. All statistical analysis was performed with GraphPad Prism 8 software. The results were considered statistically significant if p < 0.05. *p % 0.05, **p % 0.01, ***p % 0.001.

## 3. Results

### 3.1 ILM photodisruption creates ILM pores in a specific pattern

#### 3.1.1 ILM integrity in the bovine retina

First, we evaluated whether the combination of ICG and pulsed laser irradiation was able to disrupt the bovine ILM in a specific pattern. To this end, ICG solutions with varying concentrations were applied to the ILM surface of bovine retinal explants followed by laser scanning of the samples with single 7 ns pulses (800 nm, 1.65 J/cm^2^) in a fixed scanning pattern, maintaining a pore center-to-center distance of 145 µm.

As shown in Figure 4A, immunostaining of laminin, a major component of the ILM and blood vessels, revealed that we were able of creating a specific photodisruption pattern in the ILM, demonstrating the high spatial control of ILM photodisruption. At a 0.10 mg/ml ICG concentration, the ILM surface was almost unchanged in comparison to untreated, while 0.25 mg/ml ICG resulted in regularly spaced holes in the ILM corresponding to the pulsed laser spots. With higher ICG concentrations, some regular spacing was observed but the pores were confluent, creating very large ILM perforations. Moreover, damage to the underlying retina occurred, as apparent by the absence of nuclei and adjacent structures below the ILM. A brighter appearance of the intraretinal blood vessels following photodisruption is artifactual and related to the increased antibody penetration into the retina during immunostaining. Further analysis of these images with Fiji revealed that the pore diameter (Figure 4B) and pore area (Supplementary Figure 1) correlated positively with ICG concentration, whereas the percentage of intact ILM correlated negatively with ICG concentration (Figure 4C). Pore diameter measurements were more variable at higher ICG concentrations (0.50 – 1.00 mg/ml), due irregular confluence of the pores.

**Figure 4:**
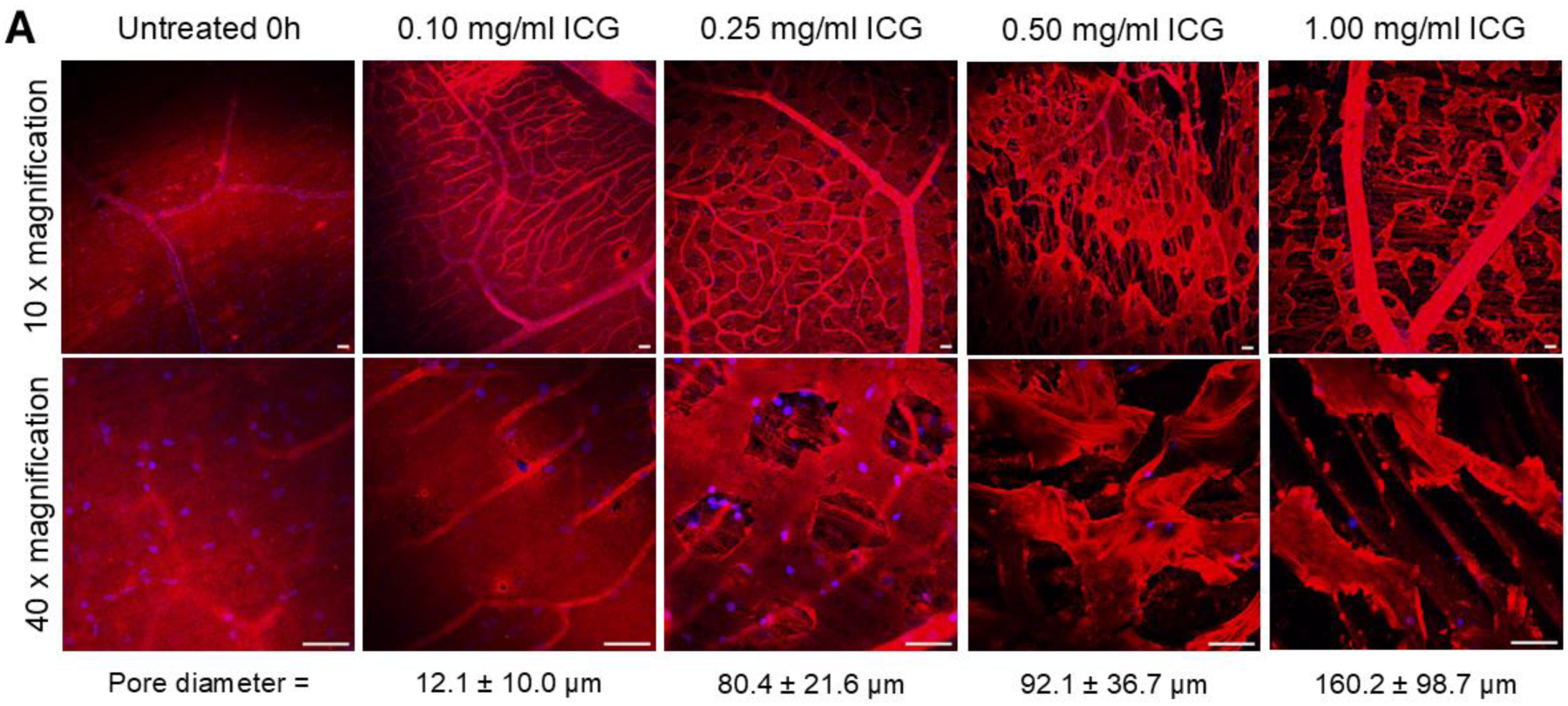
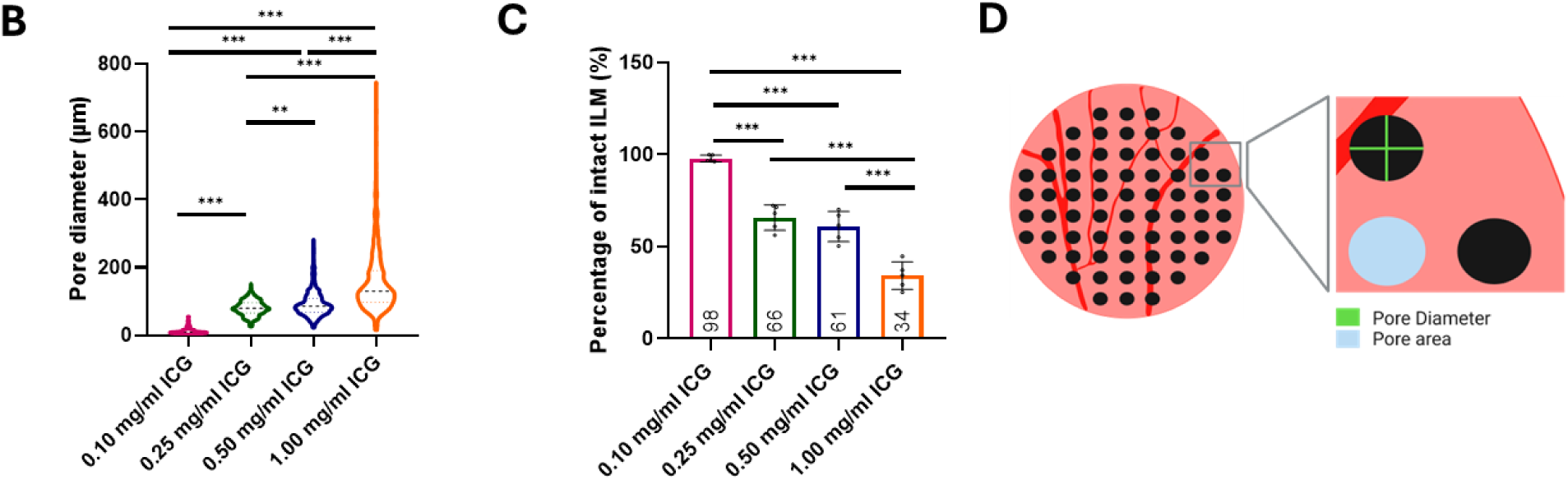
Representative images of untreated and photodisrupted bovine flatmounts. All flatmounts were immunostained for laminin (red) to investigate the ILM integrity and blood vessels. Hoechst staining (blue) was used to examine all nuclei. The confocal images reveal a distinct photodisruption pattern, highlighting the strong spatial control of photodisruption; scale bars: 50 µm (A). Quantification of the pore diameter and percentage of intact ILM for varying ICG concentrations. Increasing ICG concentrations resulted in elevated pore size (B-C). Schematic representation of pore diameter and pore area of a photodisrupted explant (D). *p ≤ 0.05, **p ≤ 0.01, ***p ≤ 0.001, NS: p > 0.05. Created with BioRender.com.

Enzymatic digestion of the ILM has been successfully applied before to disrupt the ILM and boost engraftment of stem cell-derived RGCs within the murine retina (17,30,37,38). Therefore, we selected the ILM digesting enzyme, collagenase as a positive control throughout all experiments. To identify the influence of collagenase on human and bovine ILM morphology, we applied 5 µl of collagenase with activities ranging 50 U/ml to 200 U/ml per 5 mm explant.

As expected, the morphology of the ILM following collagenase treatment differed markedly from what was observed after photodisruption (Figures 4-5). Predictably, higher concentrations, and consequently increased collagenase activity, resulted in more extensive disruption of the ILM. Besides the ILM, collagen also represents one of the main components of the retinal blood vessels, and expectedly higher collagenase concentrations also induced structural damage to the blood vessels (Figure 5). Similarly to the confocal images of photodisruption, brighter labeling of blood vessels was observed due to increased antibody penetration during immunostaining. However, due to enzymatic digestion, the integrity of these blood vessels was also compromised.

**Figure 5:**
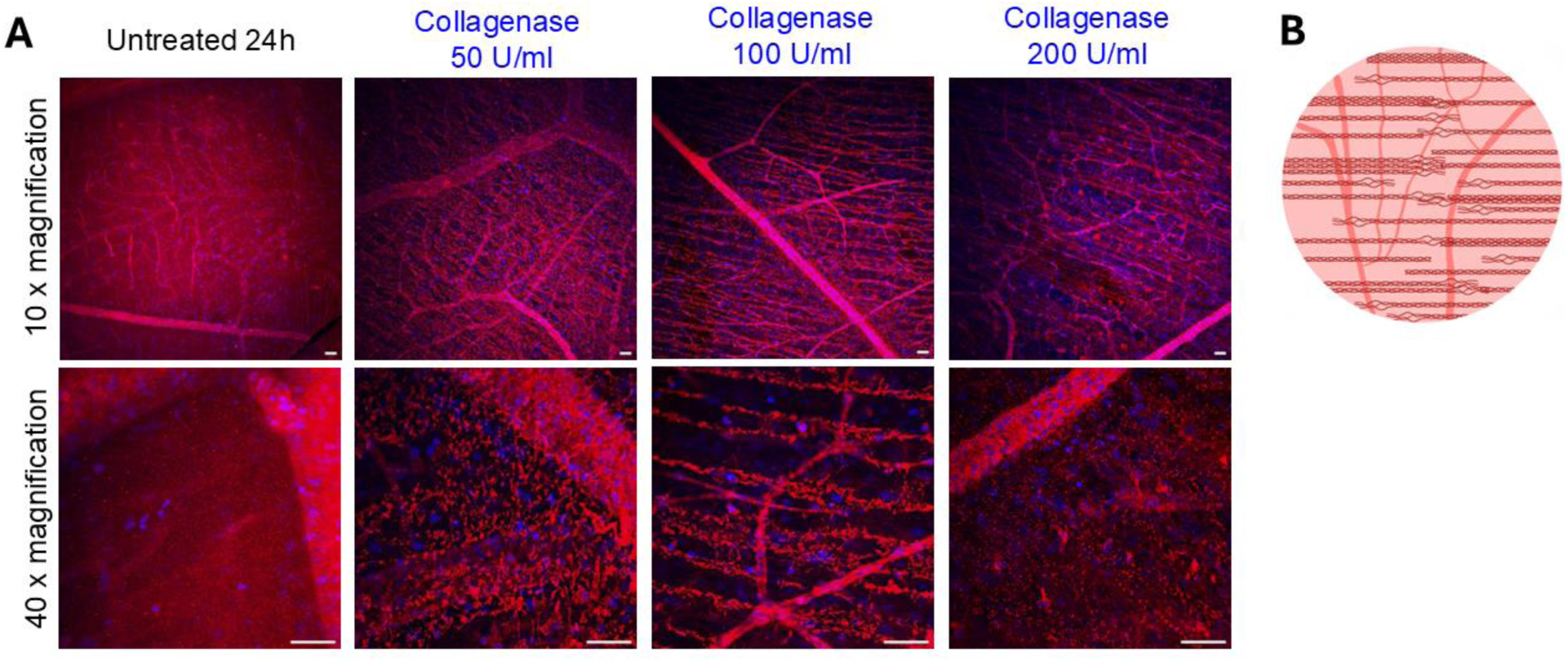
Representative images of untreated and collagenase treated bovine flatmounts. Flatmounts were immunostained for laminin (red) to investigate the ILM integrity and blood vessels. Hoechst staining (blue) was used to examine all nuclei. Increasing collagenase activity resulted in more ILM digestion and damaged blood vessels; scale bars: 50 µm (A). Schematic representation of a collagenase treated explant (B). Created with BioRender.com.

#### 3.1.2 ILM integrity of the human retina

The ILM is known to be a highly complex basement membrane with interspecies variability in thickness, morphology, composition and structure. The thickness of the bovine ILM is estimated between 100 nm and 120 nm, comparable to the ILM thickness of a human fetus (45,59). The human ILM, however, thickens with age, ranging up to a few microns in elderly patients (16,45,59). Furthermore, we previously observed that the human ILM has a much stronger affinity for ICG compared to that of bovines (45). In view of these interspecies differences and to realistically estimate the clinical potential of photodisruption, we evaluated the impact of photodisruption on human ILM integrity. Importantly, confocal microscopy on human retinal flatmounts revealed successful creation of regularly spaces photodisruption patterns in the thick human ILM of 5 different donors when combining our laser pattern with varying concentrations of ICG (Figure 6A). We did observe a more pronounced effect in the younger donor (65y), where pores seem to traverse the entire membrane, likely due to a thinner ILM, compared to the older donor (76y) where pores seem to remain more superficial, presumably due to a thicker ILM (59). Additionally, the human ILM exhibited varying morphology even in the untreated flatmounts, which was also seen when assessing pore morphology after photodisruption. Pore diameters and the percentage of intact ILM were determined based on confocal images of all 5 donors. Similarly to the bovine explants, we observed that increasing ICG concentrations resulted in larger pore diameters and lower percentages of intact ILM (Figures 6B and 6C).

**Figure 6:**
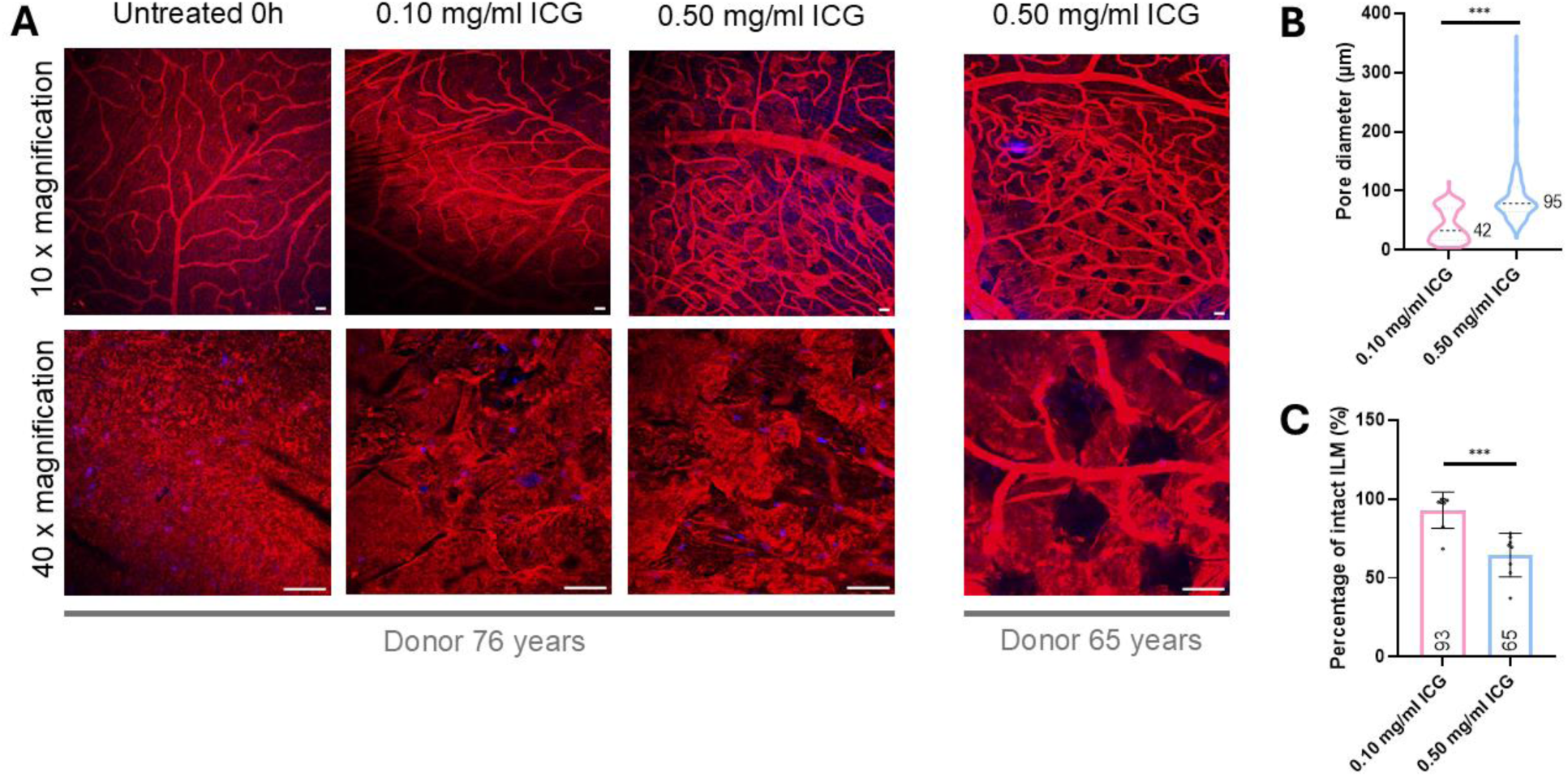
Representative confocal images of untreated and photodisrupted human flatmounts. Flatmounts were immunostained for laminin (red) to investigate the ILM integrity and blood vessels. Hoechst staining (blue) was used to examine all nuclei (A). Photodisruption reveals a distinct pattern, seen by black holes that appear in the intact (red) ILM; scale bars: 50 µm (A). Pore diameter and percentage of intact ILM were compared for varying ICG concentrations in human flatmounts of 5 donors (aged 65-78y) (B-C). Group means were compared using an unpaired t-test. *p ≤ 0.05, **p ≤ 0.01, ***p ≤ 0.001, NS: p > 0.05.

Interestingly, we obtained substantially larger pores in the thick human ILM (42.4 ± 28.2 µm), compared to the thinner bovine ILM (12.1 ± 10.0 µm) under the same conditions of 0.10 mg/ml ICG (Figures 4B and 6B), which we attribute to the human ILM’s higher affinity of ICG. It should be noted that under these conditions, we still find a high percentage of intact ILM (93 ± 11%) (Figure 6C), even though high-magnification images of the human explants treated with 0.1 mg/ml ICG demonstrate disruption of the ILM. When superficial craters were created, however, these lead to an overestimation of the effective percentage of ‘intact’ ILM. Nonetheless, when using a higher concentration of 0.5 mg/ml, no large differences in ILM intactness occurred between the bovine and human explants (Figures 4 and 6).

Next, we investigated the effect of enzymatic digestion (i.e. collagenase) on the morphology of the human ILM. Surprisingly, confocal microscopy of the human ILM did not reveal clear signs of digestion, as ILM integrity seemed comparable between the control and collagenase treated flatmounts (Figure 7), even for the 49-year-old donor who is relatively young and thus assumably has a thinner ILM (59). This is in stark contrast to the ILM digestion that we observed after collagenase treatment of bovine flatmounts (Figure 5), demonstrating clear inter-species differences in enzymatic digestion of the human versus bovine ILM.

**Figure 7:**
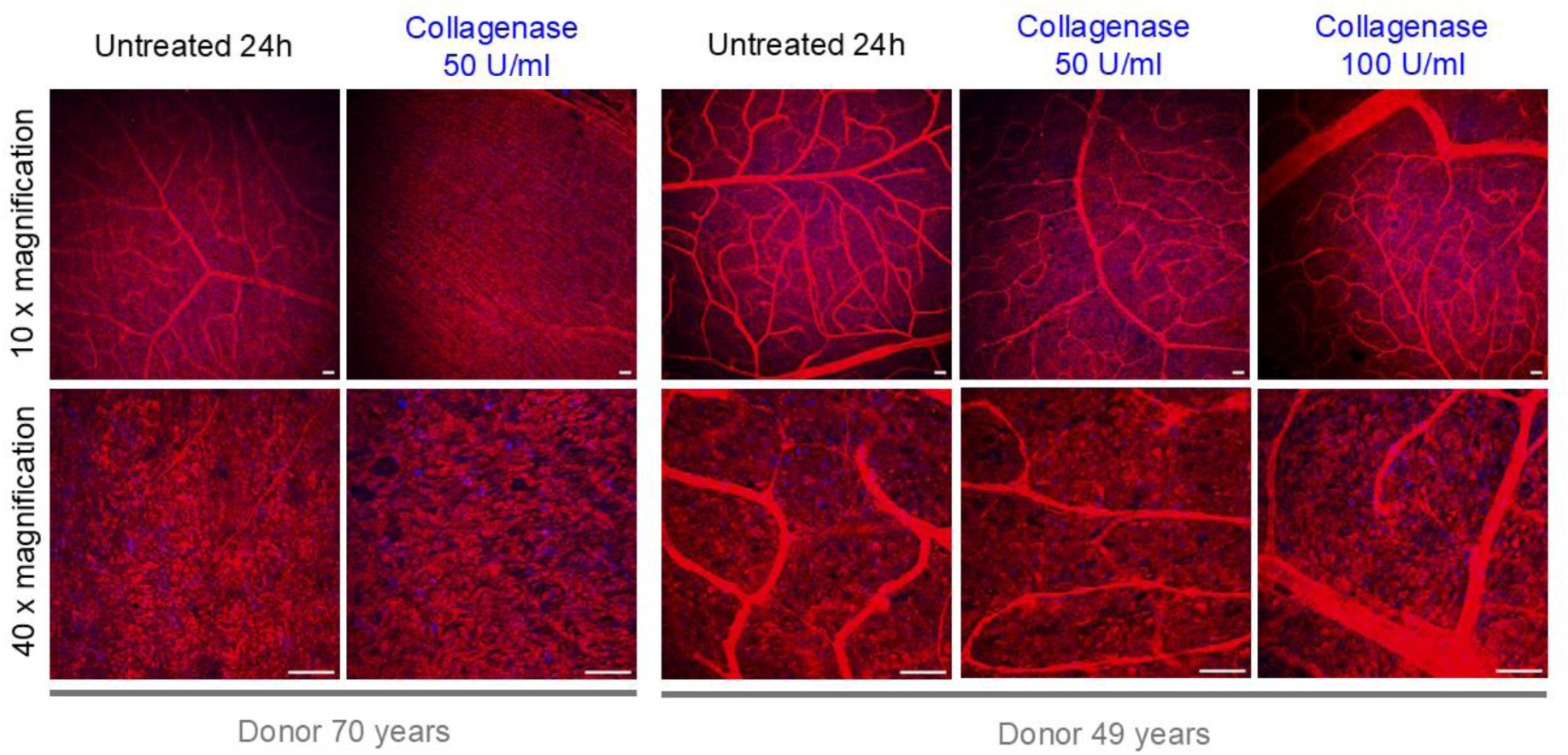
Representative images of untreated and collagenase treated human flatmounts. Flatmounts were immunostained for laminin (red) to investigate the ILM integrity and blood vessels. Hoechst staining (blue) was used to examine the nuclei. No noticeable digesting effect of collagenase was observed in the human flatmounts; scale bars: 50 µm.

### 3.2 Photodisruption augments RGC survival and neurite localization

#### 3.2.1 Donor RGC survival and density increases with photodisruption

To our knowledge, the size of human RGC in suspension is not yet known, however, RGC soma diameters cultured *in vitro* or *in vivo* range from 10 to 30 µm (49,60,61), as is comparable to the diameter of other cell types in suspension (62,63). To ensure optimal RGC migration through the created pores, while maintaining sufficient ILM integrity, we selected the ICG concentration of 0.25 mg/ml (average pore size: 80 ± 22 µm) for use in RGC transplantation experiments. We hypothesized that this pore size would allow efficient passage into the retina, while at the same time preserving a large fraction of the ILM (66 ± 6%).

To explore the impact of ILM photodisruption and collagenase treatment on RGC engraftment, hiPSC-derived RGCs expressing TdTomato (20.000 or 33.000 cells per 5 mm explant) were cocultured for 7 days with the pre-treated explants. On day 7, samples were fixed and stained for human nuclear antigen to identify the RGCs, allowing to quantify the number of RGCs per explant using confocal microscopy. Using hiPSC-RGCs from the same vial as the transplantation, a separate culture of 8000 RGCs per well was cultured in a 96-well plate for 7 days as control.

Confocal images (10x objective) of retinal explants 7 days post-transplantation, revealed RGC accumulation in regions adjacent to blood vessels, likely due to the thinner ILM in these regions, especially in the untreated explants (Figure 8A, yellow dots) (64). Interestingly, treated explants exhibited more transplanted RGCs and greater dispersion compared to untreated explants. However, RGCs on ILM-disrupted explants were still more clustered than RGCs cultured in 96-well plates. These observations were confirmed in density heat maps where the extent of RGC clustering is observed from the yellow to red color, while blue to green regions represent more dispersed cells (Figure 8B). These findings were also corroborated by various spatial metrics, which will be discussed in more detail later (section 3.2.2).

**Figure 8:**
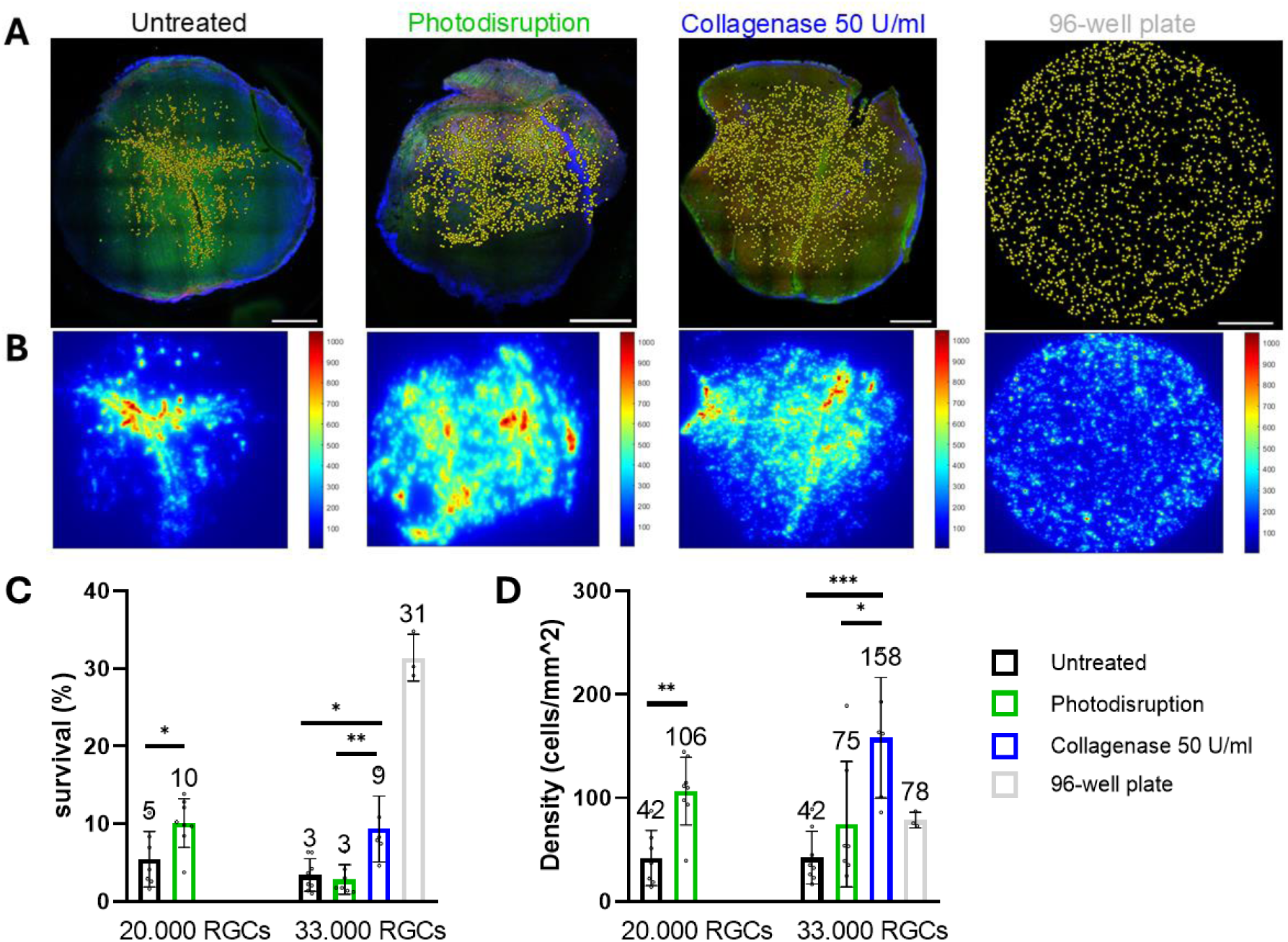
Representative confocal images of untreated, photodisrupted, and collagenase treated bovine flatmounts, 7 days after transplantation with 33.000 hiPSC-derived RGCs expressing TdTomato (red). Flatmounts were stained with Human Nuclei (green) to identify the donor RGCs and Hoechst (blue) to visualize all nuclei; scale bars: 1 mm (A). From the coordinates of the donor RGCs, density heat maps were generated using MATLAB, where red colored regions point to clustered RGCs whereas blue colored regions indicate cells that are more spread out (B). Supplementary Figure 2 contains representative confocal images and heatmaps of the transplantation involving 20,000 donor RGCs. Percent survival and density of the donor RGCs were compared for the different treatment groups and RGC starting densities (C-D). *p ≤ 0.05, **p ≤ 0.01, ***p ≤ 0.001, NS: p > 0.05.

By counting the number of integrated donor cells in the intact explants, RGC survival was compared for two different RGC starting densities (i.e. 20.000 and 33.000 RGCs per 5 mm explant), as the initial density is known to influence the survival rate of transplanted cells (7,30) (Figure 8C). Overall, a higher number of transplanted RGCs resulted in a lower survival rate. Interestingly, RGC survival in photodisrupted explants was significantly augmented, with an approximately 2-fold increase in survival rate (i.e. 10.09 ± 2.94% versus 5.42 ± 3.32% (p < 0.05) for untreated, after administration of 20.000 donor RGCs). Collagenase had a similar effect in the more densely transplanted explants wherein RGC survival increased from 3.40 ± 1.97% to 9.35 ± 3.87% (p <0.05) (Figure 8C). As expected, the highest survival rate (31.40 ± 2.47%) was observed for the RGCs seeded in the 96-well plate coated with laminin and PLO. Both treated groups exhibited higher RGC densities compared to the untreated group, with collagenase treatment particularly excelling in this aspect (Figure 8D).

#### 3.2.2 Donor RGC clustering decreases and regularity increases with photodisruption

Given that endogenous RGCs are organized in mosaic patterns to ensure a uniform visual field, avoiding RGC clustering is likely important for functional RGC transplantation (57,65). Reduced clustering after treatment with an ILM-digesting enzyme, has been described in literature (30,66). Interestingly, high magnification (40x or 60x objective) confocal images (Figures 9A and 10A) and density heat maps (Figure 8B) revealed more dispersed single cells and fewer clusters in the photodisruption and collagenase treated groups compared to untreated. These observations were further quantified by determining the nearest neighbor distance (NND), nearest neighbor index (NNI) and regularity index (RI).

**Figure 9:**
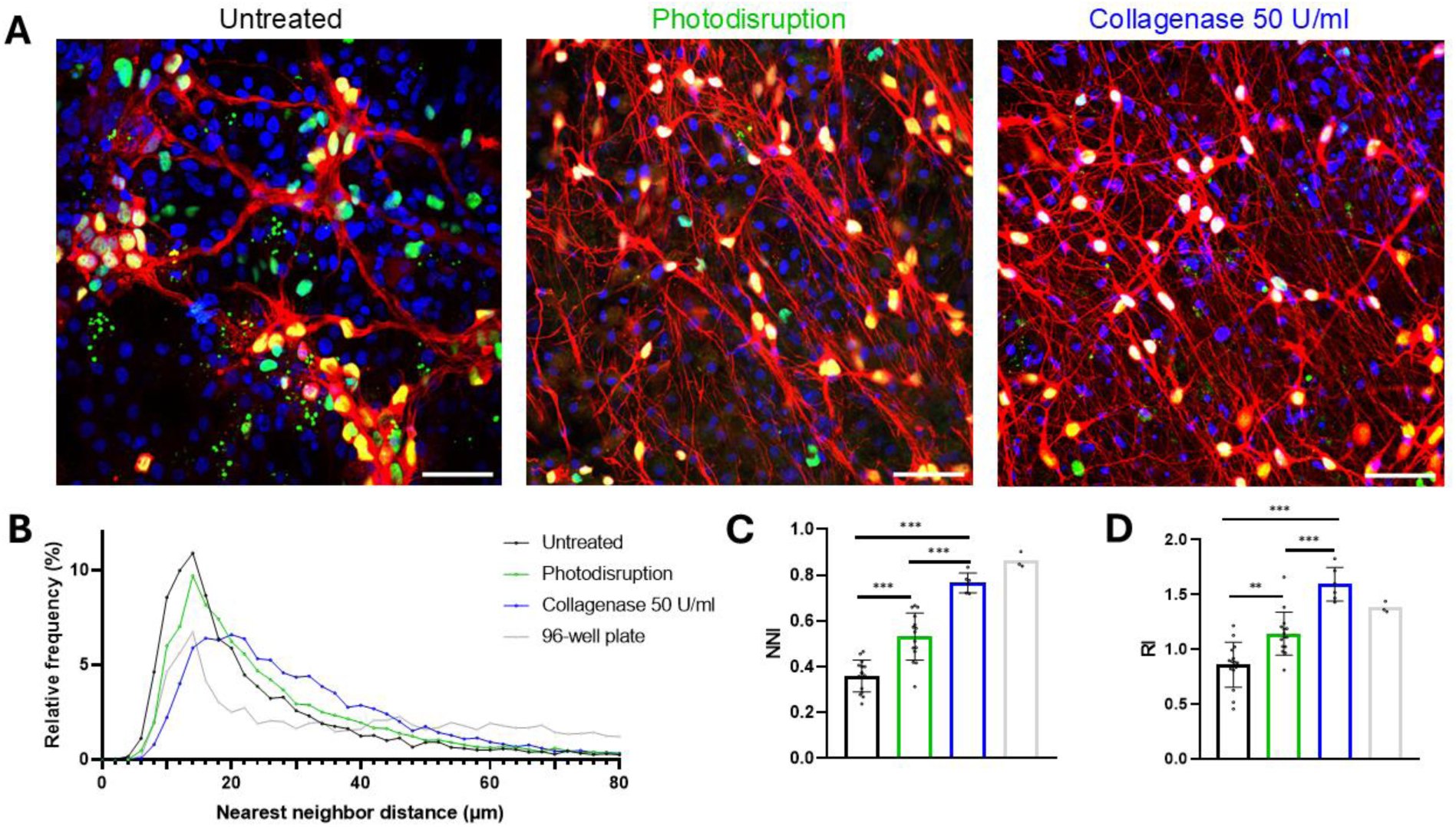
Representative confocal images (40x objective) of untreated, photodisrupted, and collagenase treated bovine flatmounts, 7 days after transplantation with hiPSC-derived RGCs expressing TdTomato (red). Flatmounts were stained with Human Nuclei (green) to identify the RGCs and Hoechst (blue) to visualize all nuclei; scale bars: 50 µm (A). Frequency distribution graph of the NNDs (B). Higher NNI values, indicating less clustering, were obtained for the treated groups compared to the untreated group (C). In addition, higher RI values were found for the treated groups, indicating a more regular RGC mosaic pattern (D). *p ≤ 0.05, **p ≤ 0.01, ***p ≤ 0.001, NS: p > 0.05.

**Figure 10:**
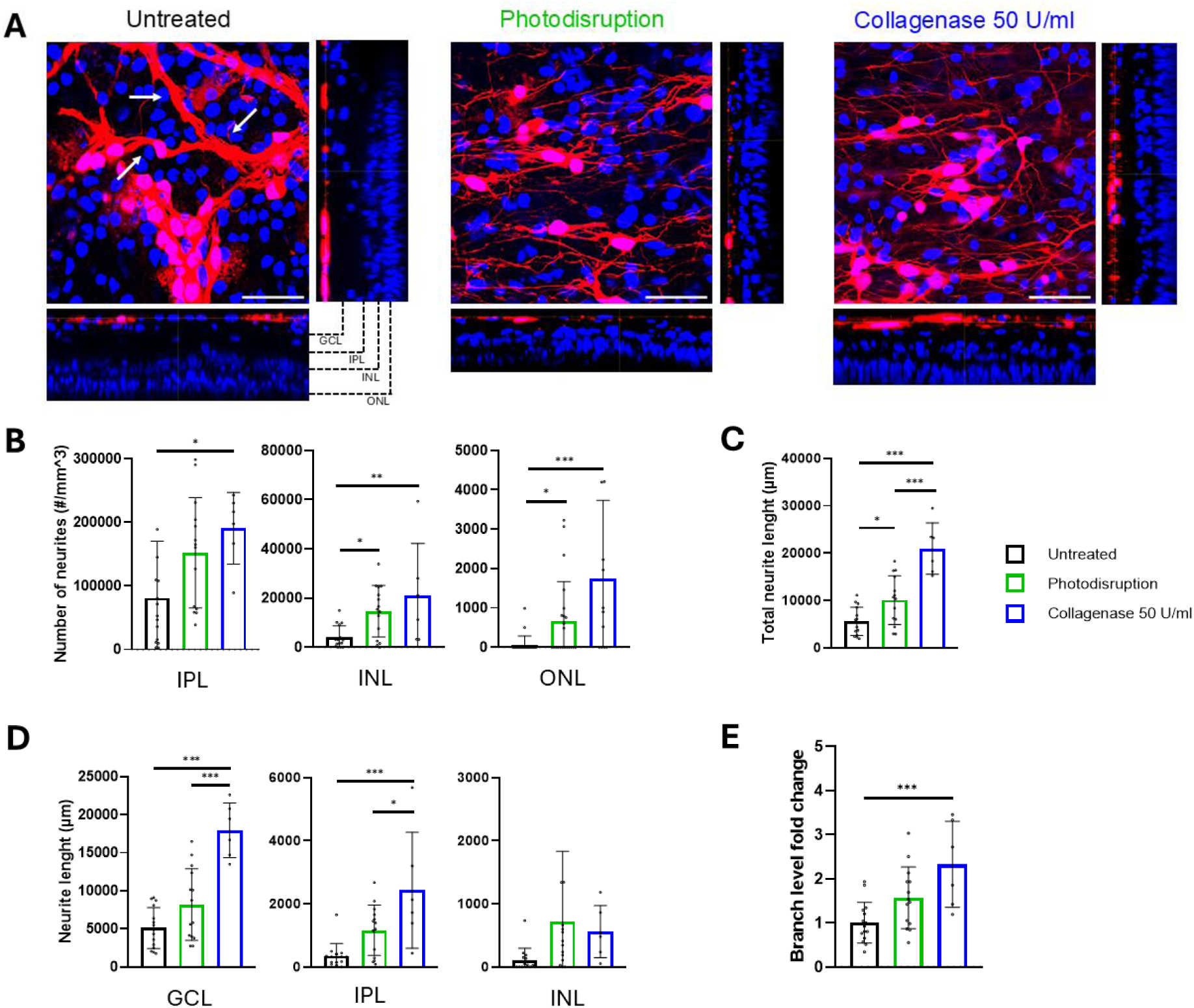
Qualitative and quantitative representation of the neurite localization of transplanted RGCs. Representative confocal images (60x objective) of untreated, photodisrupted, and collagenase treated bovine flatmounts, 7 days after transplantation with hiPSC-derived RGCs expressing TdTomato (red). Flatmounts were stained with Hoechst (blue) to visualize all nuclei. White arrows point to nerve fiber bundling; scale bars: 50 µm (A). Using Fiji, neurite localization was quantified by counting the number of neurites reaching each cell layer (B). The number of neurites per RGC reaching each cell layer is displayed in Supplementary Figure 3A. In addition, using Imaris, the total neurite length (C), neurite length per cell layer (D), and the fold change in branch level (E) were determined. Analysis with Fiji and Imaris revealed that our light-based technique and collagenase treatment were both beneficial for the RGCs neurite localization. *p ≤ 0.05, **p ≤ 0.01, ***p ≤ 0.001, NS: p > 0.05.

The NND reflects the spatial distribution of RGCs, by quantifying the distance between a reference cell and its closest neighboring transplanted RGC (30,56). Interestingly, we noted that the relative frequency distribution peak of the untreated (black) and photodisruption (green) explants was located more towards the lower NND values, indicating tighter clustering, in contrast to the collagenase treated group (blue) (Figure 9B). Furthermore, the nearest neighbor index (NNI) was determined, where an NNI value below 1.0 indicates clustering, whereas higher NNI values are associated with more dispersed cells as is desired for a successful RGC transplantation (67). The NNI increased significantly from 0.36 ± 0.07 to 0.53 ± 0.10 (p < 0.0001) in photodisrupted explants and it was even more apparent in the collagenase treated explants (0.77 ± 0.04; p < 0.0001) (Figure 9C). In addition, the regularity index (RI) was used to assess the uniformity of the mosaics, where the RI of a random distribution is approximately 1.91 and higher RI values indicate increased regularity (30,54,57,58,65). Interestingly, the RI rose from 0.86 ± 0.20 to 1.14 ± 0.19 (p < 0.01) in photodisrupted retinas along with collagenase treated explants (1.59 ± 0.14; p < 0.0001) indicating increased uniformity (Figure 9D). To summarize, by applying several spatial metric tools, we observed that the ILM photodisrupted and collagenase treated explants show significantly reduced clustering and more regularity compared to the control group, confirming the observations in Figure 8.

#### 3.2.3 RGC neurite localization after ILM photodisruption

An essential part of functional integration of donor RGCs into the visual circuitry involves the sprouting of dendrites into the inner plexiform layer (IPL) to initiate synaptogenesis with bipolar and amacrine cells (27,61). Therefore, the number of neurites reaching each cell layer was quantified in high magnification (60x objective) Z-stacks using Fiji.

As shown in Figure 9A and Figure 10A, significant nerve fiber bundling was observed in the untreated group, likely due to cell clustering. In the treated groups, on the other hand, donor RGC neurites were more homogenously spread. The number of donor RGC neurites reaching each retinal layer was calculated, distinguishing layers based on differences in Hoechst intensity using orthogonal viewing or 3D rendering in Fiji, as shown in Figure 10A and Supplementary movie 4. This analysis revealed that more neurites extended deeper into the retina for both ILM disruption treatment groups. Both ILM photodisruption and collagenase treatment were associated with more neurites in the inner plexiform layer (IPL), along with the inner nuclear layer (INL) and the outer nuclear layer (ONL) (Figure 10B and Supplementary Figure 3A).

To further substantiate this key finding, neurite localization was determined semi-automatically using Imaris-filament tracer by a second independent, masked observer. Quantitative analysis of the traced neurites revealed that neurite length and the dendritic arbors in each cell layer increased following each ILM-disrupting treatment (Figure 10D and Supplementary Figure 3C), confirming our results previously determined in Fiji. Furthermore, the total neurite length (Figure 10C), the neurite length per RGC (Supplementary Figure 3B), and the dendritic arbor area per RGC (Supplementary Figure 3D) increased after photodisruption and collagenase treatment. Finally, the fold change in branch level increased as well after treatment from 1.00 ± 0.44 to 1.57 ± 0.67 (ns, p = 0.070) for photodisruption and to 2.33 ± 0.89 (p < 0.001) for collagenase treatment (Figure 10E). Taken together, photodisruption and collagenase treatment promoted RGCs neurite formation, leading to increased neurite outgrowth and extension, with the most pronounced effects observed in the collagenase-treated group.

### 3.3 Retinal health and gliotic response after photodisruption

Examining potential retinal toxicity is a key metric in assessing the safety of this potential translational approach of ILM photodisruption. Therefore, the number of endogenous RGCs immunostained with RBPMS (green) were manually counted in bovine flatmounts immediately after dissection (0h) or after 24h, for photodisruption and collagenase treatment, respectively. At the 0h timepoint, a statistically significant decrease in endogenous RGC survival was observed for photodisrupted explants (274 ± 16 RGCs/mm^2^) compared to untreated explants (554 ± 54 RGCs/mm^2^; p < 0.05) (Figure 11 A-B). At the 24h timepoint, the number of endogenous RGCs in untreated flatmounts was more than halved (from 554 ± 54 RGCs/mm^2^ to 203 ± 6 RGCs/mm^2^; p < 0.01) compared to the 0h control (Figure 11A-B). However, no additional loss of RGCs due to collagenase treatment was observed (Figure 11A-B). Next, we evaluated retinal thinning of the explants 7 days after transplantation by comparing the distance of the different cell layers from the ILM using Fiji (i.e. ILM to IPL, ILM to INL, ILM to ONL). As shown in Figure 11C, no significant differences were observed, indicating no substantial retinal thinning induced by either treatment, above what occurred normally during the culture period.

**Figure 11:**
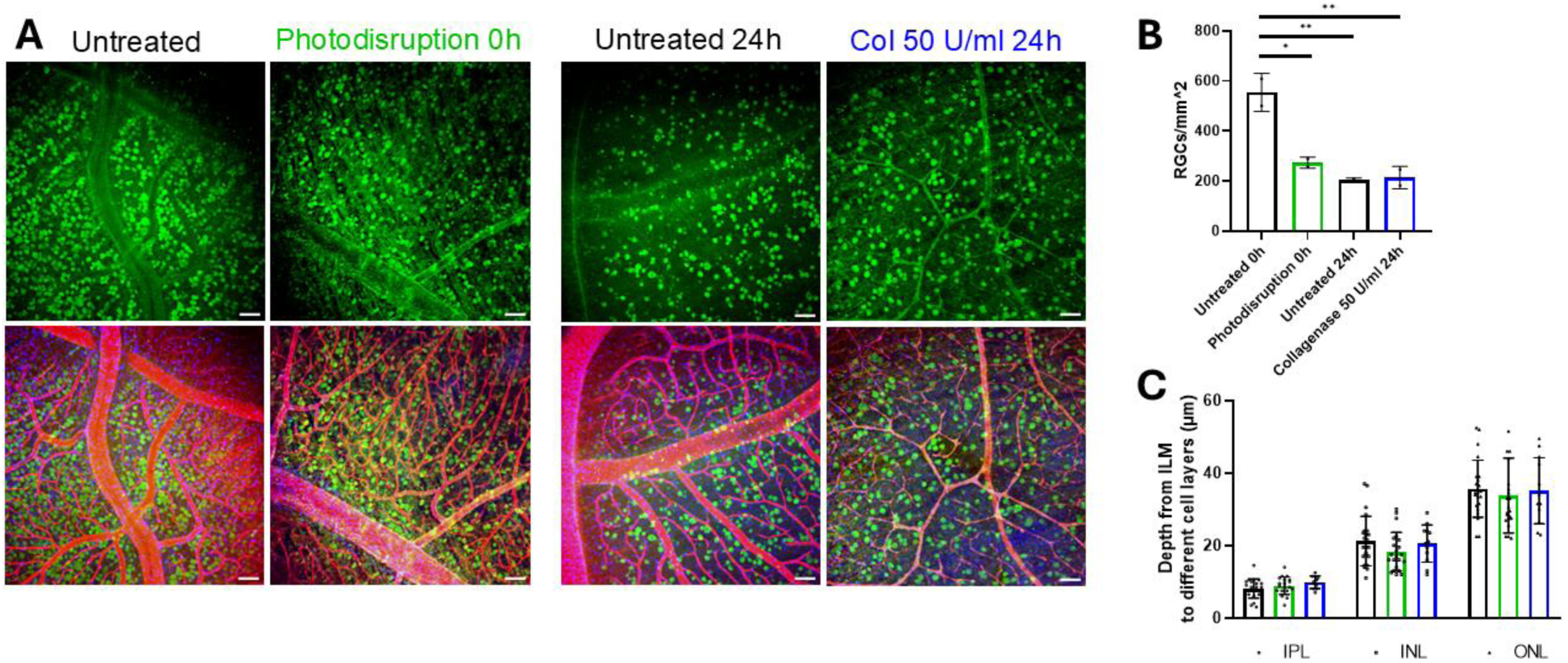
Representative detailed flatmount images (10x objective) of endogenous RGCs stained by RBPMS (green), the ILM immunostained by laminin (red), and Hoechst (blue) to stain all nuclei; scale bars: 100 µm (A). Quantitative comparison of the number of endogenous RGCs manually counted in Fiji for photodisrupted and collagenase treated explants, with their respective controls at the 0h and 24h time point (B). Quantitative image analysis using Fiji of retinal thinning 7 days after transplantation. No significant differences between groups were observed (C). *p ≤ 0.05, **p ≤ 0.01, ***p ≤ 0.001, NS: p > 0.05.

Another important factor for successful transplantation is the gliotic response mediated by the host (i.e. bovine explant), as this can create either a protective or neurotoxic environment for donor RGC integration and survival. Indeed, the success of transplantation may be contingent upon the response orchestrated by the resident innate immune cells within the explants, including Müller glia, astrocytes, and microglia. While neuroinflammation may potentially be exacerbated by the xenogeneic nature of the human iPSC-derived RGCs and immunosuppression was not employed in our experiments, it should be noted that there was no peripheral circulation in these cultures and therefore no potential for an adaptive immune response. Nonetheless, explantation and culturing of the explants is known to induce gliosis. To investigate the latter, we stained for glial fibrillary acidic protein (GFAP) (green), a well-established marker of retinal stress in Müller glia and astrocytes. Following the 7-day coculture with RGCs, confocal microscopy revealed a similar level of GFAP expression for both untreated and treated bovine explants (Figure 12A), indicating our treatments did not evoke an additional gliotic response.

**Figure 12:**
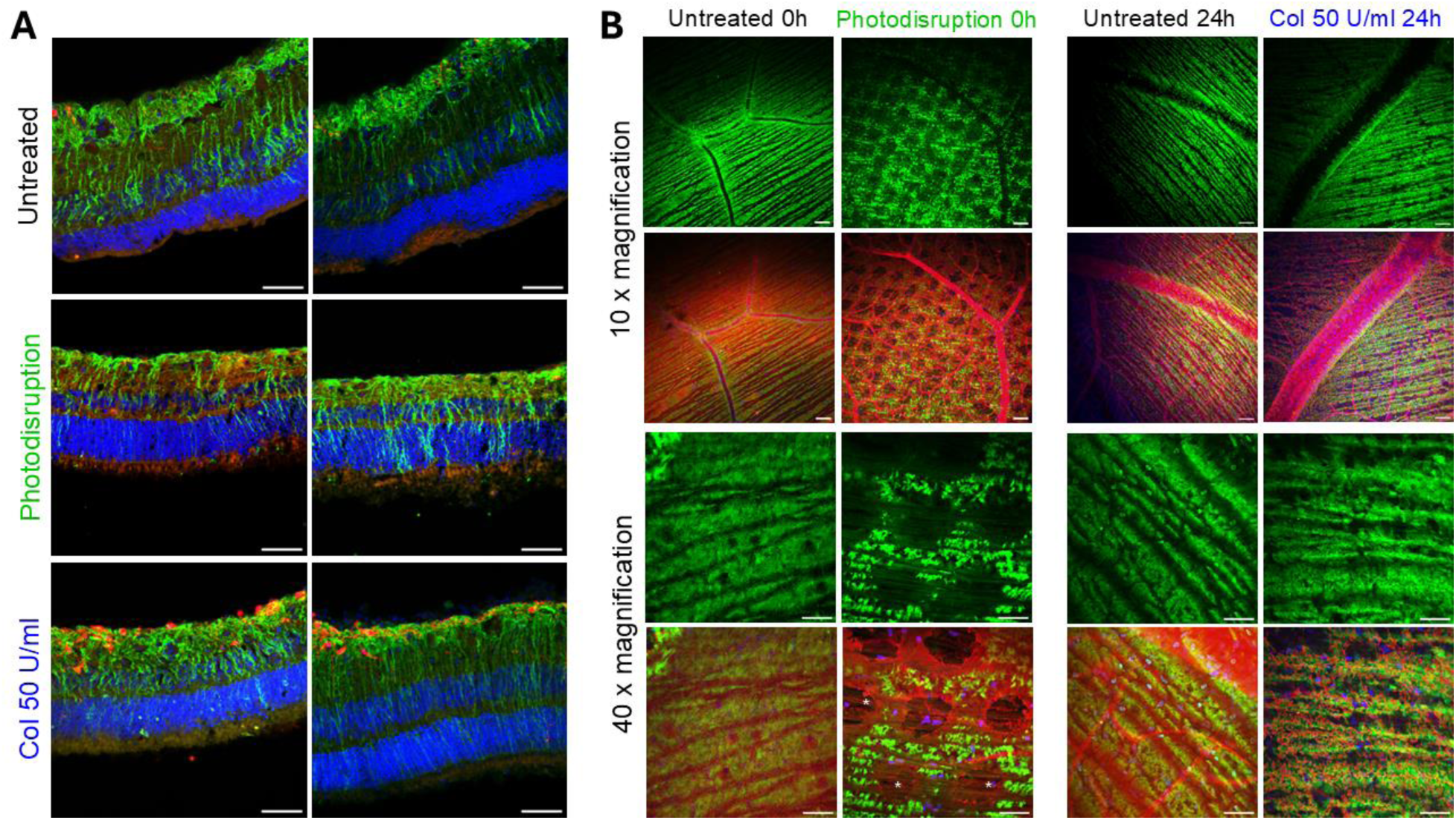
Representative images of the gliotic response of bovine retinal explants, 7 days after transplantation with hiPSC-derived RGCs expressing TdTomato (red). Cryosections were stained for Hoechst (blue) to visualize nuclei and GFAP (green) to visualize glial reactivity in Müller glia and astrocytes; (40x objective) scale bars: 50 µm (A). In another experiment without RGC transplantation, flatmounts were stained with GS to visualize Müller glia (green), laminin (red) to examine the ILM, and Hoechst (blue) to investigate nuclei. The photodisruption pattern was clearly recognizable in both the Müller cell staining and in the ILM staining, whereas the Müller endfeet damage was colocalized with the ILM damage after collagenase treatment. Intact nerve fibers were observed after photodisruption, as indicated by the white stars; scale bars: 100 µm for 10x magnification images and 50 µm for 40x magnification images (B).

Additionally, we assessed the morphology of the Müller glia cells, considering their endfeet are located just below the ILM, by immunostaining for glutamine synthetase (GS) (green) in bovine flatmounts without RGC coculture (Figure 12B). In the untreated explants, Müller cells appeared homogenous, and the ILM was intact. In the photodisrupted explants, on the other hand, the individual pores created by the laser pattern were clearly recognizable in both the Müller cell staining and in the ILM staining; indicating that the laser treatment induced localized damage to the Müller cell endfeet (Figure 12B). Importantly, in high magnification images, we observed that the nerve fiber layer appeared to remain intact, and moreover, normal appearing cell nuclei were present at the locations of photodisruption. In the explants subjected to collagenase treatment, the Müller cell endfeet damage was colocalized with the ILM disruptions. Although less apparent, collagenase treatment led to a more faded appearance of Müller endfeet as opposed to untreated explants. Furthermore, the degree of GS damage increased with the collagenase activity (data not shown).

In summary, photodisruption showed some adverse effects, such as a reduced number of endogenous bovine RGCs and damage to Müller cell end feet, suggesting a potential negative impact on retinal health. However, these effects appeared relatively mild, as no significant retinal thinning was observed. In addition, Müller glia and astrocytes reactivity remained consistent with expectations after 8 days in culture. Nevertheless, further research is needed to fully assess the safety of this light-based therapy.

## 4. Discussion

This study explored the potential of ICG-mediated photodisruption as a strategy to selectively perforate the ILM in a predetermined pattern, to enhance its permeability and improve RGC delivery and subsequent engraftment. By creating a distinct pattern of pores within the ILM, we hypothesized that we could establish entryways for the RGCs while also preserving a substantial amount of the membrane’s integrity to support the development of the newly transplanted RGCs. To assess effectiveness, we compared photodisruption to collagenase treatment which is a well-established strategy to enzymatically disrupt the ILM. This discussion first addresses collagenase results, followed by an analysis of the photodisruption outcomes.

### Enzymatic digestion of the ILM

Enzymatic digestion has been widely employed to modify the ILM, with various enzymes tested for several therapeutic applications, including AAV gene therapy and cell therapy (30,34,35,37,38,68). Digesting the ILM has demonstrated efficacy in enhancing delivery across multiple animal models, including mice, rats and non-human primates. However, this approach also has a significant drawback: the risk of bleeding. This is unsurprising, as collagen—a primary target of ILM digestion—is also a key structural component of blood vessel basement membranes (69,70). Previous research has highlighted the narrow therapeutic window for these ILM-digesting enzymes, emphasizing the delicate balance required for their use (35,37). Indeed, our study corroborated these findings, showing that collagenase treatment effectively broke down the bovine ILM as well as the blood vessels (Figure 5). Notably, our study is the first to examine the morphology of the human ILM following collagenase digestion where we found that concentrations that were highly effective in breaking down the bovine ILM showed no apparent effect on the human retina (Figure 7). Importantly, the range of human tissue donor age in this study aligns with that of individuals typically affected by (chronic) glaucoma. The inability of collagenase to disrupt the human ILM in these samples thus raises significant doubts about its clinical feasibility for glaucoma patients. Moving forward, we aim to repeat these studies using higher collagenase activities, assuming these might be effective in disrupting the thick human ILM, although this carries the risk of exacerbating toxicity and intraocular hemorrhage.

In the next phase of the study, we looked into the impact of enzymatic digestion on RGC delivery and survival in bovine flatmounts. Our survival rates of transplanted RGCs were comparable to those reported in literature (5 up to 25%) for *ex vivo* experimentation (6,7,9,30,71,72). In our study, two different transplantation densities (i.e. 20.000 RGCs or 33.000 RGCs per explant) were compared. Interestingly, the lower starting density resulted in a higher percental survival rate (Figure 8). This counterintuitive outcome was also reported by Venugopalan *et al.* (7) and may be attributed to factors such as nutrient depletion, increased gliotic response, and a potential reverse U-shaped relationship between transplanted cell quantity and survival (11,30). It must be noted that variations in outcomes, including our relatively low survival rate, may be explained by differences in RGC origin (hESC or iPSC), recipient species, animal model type, and duration of RGC coculture.

Importantly, ILM digestion with collagenase significantly increased the percent survival and density of donor RGCs in our bovine explants (Figure 8). This confirms our previous work where we revealed that RGC survival was increased after applying the digestive enzyme pronase (37,38). Next, we investigated RGC clustering since this might prevent donor RGCs to carefully organize themselves in mosaic patterns necessary to form a uniform visual field. Previous research indeed revealed that the average NND of transplanted RGCs in controls significantly increased after digestion with pronase or collagenase, suggesting that enzymatic treatment reduces clustering (30,66). Analogously, in our bovine retinas, more dispersed cells and less clustering were visually observed after ILM digestion, which again suggests that interactions between RGCs and the ILM promote clustering (Figure 9) (30). To quantify our qualitative observations, we employed several spatial analytic tools. We first examined the NNI values of the donor RGCs, which more than doubled in collagenase-treated explants compared to those with an intact ILM, implying a substantial reduction in cell clustering (Figure 9). Interestingly, our findings align with previous studies, including our own, which demonstrated that NNI values of human stem cell-derived RGCs transplanted into murine retinas ranged from 0.40 to 0.80 (9,30). For comparison, we calculated the NNI of endogenous RGCs in our bovine explants, which was 1.1 ± 0.1 - remarkably higher than the NNI values observed for transplanted RGCs in both murine and bovine explants. Similarly, endogenous murine RGCs exhibit NNI values of approximately 1.3 ± 0.1, comparable to the endogenous NNI values in bovines (30,67). Nevertheless, all NNI values following transplantation remained below the threshold of 1.0, indicating that clustering did persist across all groups (56,67) and complete cell dispersion was not achieved. Next, the RI was calculated to examine the regularity of the neuronal mosaics. As a reference, we characterized the RI of endogenous RGCs in the bovine retina, finding it to be 2.9 ± 0.5. This value is slightly lower than the reported RI range of 3 to 6 in mice, highlighting species-specific differences in retinal organization (54,57). Since cell density, cell type, cell size and cell death can all have a profound impact on the RI, it is difficult to compare the RI between different studies (30,54,57,58,65). Therefore, RI values were only compared within our own experiments were we discovered that the RI rose significantly after collagenase treatment suggesting an increase in regularity (Figure 9). To summarize, by applying several metric tools, we observed that collagenase treatment resulted in significantly less clustering and more uniformity which is desirable for a successful RGC engraftment.

The success of RGC transplantation is heavily influenced by the functional integration of donor cells into the visual circuitry, which requires neurite sprouting toward IPL and the formation of synaptic connections (61). Preferably, as in the collagenase treated bovine flatmounts, neurites without thick nerve fiber bundles are obtained to be able to cover the visual field uniformly (Figures 9-10). Importantly, after ILM digestion, the transplanted RGCs developed significantly more neurites that extended deeper into the retina to the IPL, along with the INL and ONL (Figure 10 and Supplementary Figure 3). Indeed, analogously to what we previously observed in mice, ILM digestion resulted in enhanced neurite integration (i.e. significantly longer neurite length and more neurite segments in IPL, INL and ONL) (30,37,38,66). Our observations in bovine flatmounts, quantified using Fiji software, were further validated through neurite tracing with Imaris, where an increase in neurites length per cell layer, neurite area, total neurite length and neurite area per RGC following collagenase treatment was confirmed (Figures 10 and Supplementary Figure 4). In addition, the branch level doubled, suggesting enhanced complexity in RGC neurite branching and an improved integration following enzymatic digestion (Figures 10). Interestingly, donor RGC neurites did not exclusively target the IPL but also extended into deeper retinal layers such as the INL and ONL. This phenomenon has been reported multiple times in studies investigating RGC transplantation in mice (9,30,71,72). Such extensions beyond the IPL, into deeper layers of the retina are undesirable and may be artifacts of explant culture, as they are observed to a much lesser extent *in vivo* (38). In sum, collagenase treatment was beneficial for RGC neurite localization as the neurons developed more complex neurites that extended further into the retina.

A critical factor influencing the success of RGC transplantation is the gliotic response of the host tissue, mediated by Müller glia, astrocytes, and microglia (14). Treatment with the gliotoxic agent alpha-aminoadipic acid which suppresses glial reactivity led to a 50-fold increase in stem cell migration into rat retinal explants, highlighting the importance of the gliotic response on transplantations (35). Interestingly, GFAP staining - an established marker of retinal stress in astrocytes and Müller cells - appeared similar across all experimental groups (Figure 12). However, increases in GFAP expression in response to our ILM treatments might be masked by the gliosis induced through explantation and culturing of the explants. These findings confirm our previous observations in murine explants, where GFAP expression remained similar following pronase treatment (30).

Looking further into collagenase’s impact on retinal health, we observed no significant retinal thinning nor additional endogenous RGC death beyond that attributable to the culturing process (Figure 11). However, a striking observation was the near 50% reduction in endogenous bovine RGCs after 24 hours of culture, even without treatment (Figure 11). This gradual RGC loss in explants is well-documented in the literature. In fact, we have proposed that *ex-vivo* explants may serve as models for optic neuropathies like glaucoma due to this phenomenon (73,74). Additionally, collagenase treatment appeared to alter the morphology of the Müller cell endfeet, as visualized by glutamine synthetase (GS) immunostaining (Figure 12). The endfeet appeared less defined and more diffuse compared to untreated explant. This endfeet damage was furthermore colocalized with the ILM disruption and increased with higher enzyme activities (data not shown).

To conclude, this study shows for the first time that collagenase-induced digestion of the bovine ILM resulted in enhanced ILM permeability, leading to an improved RGC engraftment (i.e. increased donor RGC survival, density, regularity, neurite localization, and branching). These results confirm our previously published results in murine explants, where we saw reduced RGC clustering and increased retinal neurite ingrowth after enzymatic ILM digestion (30,37). Despite its effectiveness in enhancing RGC transplantation, the enzyme-induced damage to retinal blood vessels and underlying tissue raises serious concerns on retinal health, significantly limiting its potential for clinical translation (13,30,34,35).

### ILM photodisruption

We explored the potential of ICG-mediated photodisruption as a strategy to selectively perforate the ILM in a predetermined pattern. Interestingly, microscopy images of photodisrupted flatmounts revealed the distinct and tunable pattern that was created when combining pulsed laser irradiation with varying ICG concentrations. As expected, increasing concentrations of the photothermal agent ICG, led to statistically significant elevated pore diameters and pore area (Figure 4 and Supplementary Figure 1), demonstrating the tunability of our light-based approach. Furthermore, we successfully created a distinct photodisruption pattern in the thick and complex human ILM of 5 different donors (Figure 6), which strongly contrasts with enzyme application, where we observed no apparent impact on the human ILM at the concentrations tested (Figure 7). As observed in bovine explants, increasing ICG concentrations resulted in larger pore diameters in human retinal tissue. However, pores appeared more superficial in explants from older donors (76 years) compared to those from younger donors (65 years), possibly due to age-related differences in ILM thickness, underscoring the importance of a tunable approach. Photodisruption offers this flexibility, allowing for adjustable settings and ICG concentrations to accommodate the diverse ILM thicknesses encountered across different patient age groups. Since a 0.25 mg/ml ICG concentration resulted in a regular pattern of pores of 80 ± 22 µm in size and left the majority of the ILM intact (66 ± 6%), we selected this condition for RGC transplantation experiments in bovine explants.

Interestingly, our light-based approach nearly doubled the survival of RGCs after administration of 20.000 RGCs, reaching 10% (Figure 8). Moreover, both starting density experiments exhibited higher RGC densities compared to the untreated group, albeit still considerably lower than those observed following collagenase treatment. Furthermore, confocal imaging revealed a more dispersed distribution of RGCs following photodisruption compared to an intact ILM, which was indeed confirmed by several spatial metrics including NNI and RI. First, the NNI increased by 47% in photodisrupted explants compared to untreated controls (from 0.36 ± 0.07 to 0.53 ± 0.10), indicating reduced clustering. Similarly, the regularity index (RI) showed a significant increase (from 0.86 ± 0.20 to 1.14 ± 0.19), reflecting enhanced spatial regularity (Figure 9). Following photodisruption, donor RGCs also developed significantly more neurites that extended deeper into the retina. While the majority of neurites were concentrated in the IPL, increases were observed across all retinal layers, consistent with the results obtained after ILM digestion. This positive observation also reflects in other parameters including neurite length, neurite area and branch level (Figure 10 and Supplementary Figure 3). Our findings indicate augmented engraftment following ILM photodisruption across multiple parameters. However, it is relevant to note that collagenase treatment yielded even more favorable outcomes, surpassing the other groups in terms of transplanted RGC survival, density, clustering, neurite localization, and branching.

Regarding the impact of our light-based approach on retinal health, several key observations were made. After 8 days of culturing, photodisrupted explants showed no retinal thinning compared to untreated samples. Furthermore, the GFAP staining appeared similar to untreated explants, indicating photodisruption did not evoke an additional gliotic response nor did it reduce it. Nevertheless, immediately following photodisruption, a significant and instant decrease in endogenous RGC survival of the treated explants was observed (Figure 11), indicating retinal toxicity. Our light-based therapy also affected Müller cells, as evidenced by the distinct photodisruption pattern visible in both the Müller cell and ILM staining (Figure 12). On the other hand, we did find intact cell nuclei at the pore sites, and underlying nerve fibers seem to remain intact (Figure 12). In the future, we aim to analyze Müller cell morphology in greater detail over time to determine whether the immediate damage to their endfeet is permanent or reversible. However, we should note that with the current laser set-up, the laser irradiation originates from beneath the retinal explant, necessitating passage through the entire retina before reaching the ICG located on top of the ILM. Future modifications to the laser configuration will allow for irradiation from the ILM side, directly targeting the ICG. This will enable the use of lower laser fluences as less energy will be lost by traversing the retina. Secondly, we will increase the pulse frequency, enabling faster processing, reducing the scanning time from approximately 15 minutes to less than 2 minutes, thereby reducing the risk of explant desiccation during the laser treatment. Notably, candidate patients for this procedure are anticipated to have minimal or no remaining RGCs, rendering potential damage to the structures beneath the ILM less significant in this context. By implementing these adjustments, we aim to mitigate toxicity in our future studies.

Importantly, while we used FDA-approved ICG as the photothermal agent in all photodisruption experiments, its use as an ILM dye has declined due to reports of retinal toxicity. Many surgeons now prefer alternative vital ILM dyes such as brilliant blue and trypan blue (75). In response, our group is exploring alternative dye options to mitigate potential retinal damage. Concurrently, we are evaluating the safety of our light-based photodisruption technology *in vivo*. Moreover, we have demonstrated that laser-induced nanobubbles can safely ablate vitreous opacities *in vivo* in rabbit models (47). These advancements suggest a promising future for our photodisruption technique in clinical applications.

Taken together, the results of this study highlight ILM photodisruption as a promising technique for enhancing RGC transplantation, offering several advantages over previously tested methods such as enzymatic digestion, ILM peeling, and mechanical disruption. ILM peeling is a surgical procedure where the ILM is removed for various diseases affecting the vitreomacular interface including macular holes and diabetic macular oedema (76) and has been proposed as an alternative method to overcome the ILM barrier to boost RGC transplantations (77). Nonetheless, it is an invasive procedure that can damage underlying structures, particularly the Müller cell end feet (40,41). Moreover, in glaucoma patients, whose retinas are already compromised, ILM peeling may pose an even greater risk of additional damage compared to its effects on healthy retinas (78,79). On top of that, ILM-RGC interactions are likely necessary for coordinated patterning of the RGC layer and correct dendritic projection, therefore complete removal of the ILM – as achieved by surgical peeling is not desirable. Another strategy is the mechanical disruption of the ILM (17), wherein cracks created in the ILM using mechanical forces led to improved neurite formation. However, our photodisruption technique offers several advantages over this mechanical approach, as it provides more precise and reproducible ILM disruption compared to the variable damage that may result from mechanical methods. In addition, mechanical disruption is a surgical method that implies a more invasive approach compared to photodisruption. In contrast, our light-based technique *in vivo* requires only a minimally invasive intravitreal injection of the photothermal dye followed by non-invasive laser treatment. This reduced invasiveness is particularly advantageous for *in vivo* applications, as it minimizes the risk of complications associated with more invasive surgical procedures.

The current data suggests that photodisruption holds great promise due to distinct advantages over alternative ILM disruption methods, even if some questions remain unanswered for now. Since collagenase treatment currently outperformed photodisruption in terms of RGC survival and integration, efforts to further optimize the therapeutic window are ongoing. Apart from adjusting the ICG concentration, the laser fluence also allows us to control the VNB size and hence the pores in the ILM. In this study, the same laser fluence (i.e. 1.65 J/cm^2^) was applied for all photodisrupted explants, though in future studies we will similarly tune the laser fluence to determine the optimal pore size. Furthermore, the laser set-up will be modified to enhance the efficiency and safety of the photodisruption process, as discussed above. Given the critical role of ILM-RGC interactions in coordinated patterning of the RGC layer and proper dendritic projection (26), future investigations will assess the impact of photodisruption on synaptogenesis, by immunostaining for mature dendrites (MAP2) (37,80), mature axons (SMI-312) (37,81), and synaptic machinery (synaptophysin and PSD-95) (37,82,83). Additionally, further research will be conducted to elucidate the migration and integration of RGCs over time following ICG-mediated photodisruption.

### Limitations

As in all studies, there are comments and limitations to our techniques and models. First, *ex vivo* retinal explants were used to assess the ILM integrity and integration of transplanted RGCs but are not completely representative of the *in vivo* eye. These *ex vivo* explants likely bias towards greater engraftment compared to intravitreal injections *in vivo*, given that the donor RGCs are maintained in direct contact with the ILM surface rather than being suspended within the vitreous. Nevertheless, *ex vivo* retina culture models enable higher throughput experiments and are in accordance with the principles of the 3Rs to replace, reduce, and refine animal research. Furthermore, it must be noted that all retinal explants were prepared from healthy non-glaucomatous bovine or human eyes. Interestingly, glaucomatous retinas represent a unique environment, given that the ILM thickness increases with age and in the presence of diseases like diabetes (59,84). Therefore, patients suffering from age-related optic neuropathies such as glaucoma may have a greater burden to RGC migration into the retina resulting in less engrafted donor neurons (71). Nevertheless, other sources suggest that glaucomatous retinas may exhibit an increased ability to accept donor RGCs compared to healthy retinas (9).

In recent years, a major concern has emerged regarding the observation of rare intercellular material transfer between donor and host cells, which has confounded the interpretation of various RGC and RPE transplantation studies (55,71). To address this issue, we implemented stringent criteria for RGC identification, counting only cells as RGCs if positive for both tdTomato and human nuclei markers. Cells that were positive for human nuclei but lacking tdTomato were classified as residual hiPSCs, and we did not observe any cells expressing tdTomato but not human nuclear antigen. Furthermore, GFAP staining of the cryosections did not elucidate any colocalization of GFAP and tdTomato (Figure 12), unlike what we previously observed in host, mouse Müller glia following ILM disruption (55). This suggests an absence of intercellular material transfer between donor hiPSC-RGCs and bovine Müller glia in our study. However, we acknowledge that this phenomenon cannot be definitively ruled out based on these observations alone. To conclusively exclude material transfer, additional methodologies would be needed such as species-specific PCR on isolated cells that are positive for both tdTomato and human nuclei markers or fluorescence in situ hybridization (FISH) on cryosections.

## Conclusion

This study provides compelling proof of concept for the use of ILM photodisruption to enhance RGC transplantation in *ex vivo* bovine retinal explants. Our comprehensive evaluation of the ILM integrity following photodisruption, as a function of varying ICG concentrations, demonstrates the technique’s high tunability and control in creating distinct photodisruption patterns in both bovine and human tissue. Furthermore, we observed augmented RGC engraftment after treatment with our light-based technique as demonstrated by a higher donor RGC survival rate, improved cell spreading and enhanced neurite formation. Concurrently, we discovered that collagenase digestion of the bovine ILM significantly enhanced RGC engraftment. However, the same enzymatic activities proved ineffective on the human ILM. This finding, coupled with collagenase’s well-documented retinal toxicity, particularly its damaging effects on blood vessels, casts significant doubt on its clinical viability for human applications. Although our current light-based technology showed some retinal toxicity, we are committed to refining our approach with future modifications focusing on optimizing the laser set-up and exploring alternative photothermal ILM dyes. In conclusion, our proof-of-concept study demonstrates that ILM photodisruption effectively addresses a critical barrier in RGC replacement. This breakthrough paves the way for advancing retinal regeneration toward clinical translation, offering new possibilities for treating irreversible blindness caused by glaucoma and other optic neuropathies.

## Funding

Acknowledgement is made to the donors of the National Glaucoma Research, a program of the BrightFocus Foundation, for support of this research (G2023002F). Additionally, this research is supported by the Research Foundation-Flanders, Belgium (FWO Vlaanderen, grant 1S19725N and 1226224N). Chloë De Clercq acknowledges support from Novistem 101071105, horizon 2021 EIC PATHFINDER. Weiran Li acknowledges the financial support from China Scholarship Council (CSC). Thomas Johnson is supported by the National Eye Institute (United States National Institutes of Health, K08EY031801, R21EY034332, and P30EY001765), Research to Prevent Blindness (New York, NY, Career Development Award and unrestricted funding to the Wilmer Eye Institute), Bright Focus Foundation (G2022005S), the Glaucoma Foundation Rajen Savjani Award, The Shelley and Allan Holt Rising Professorship, and The Zenkel Family Foundation.

We would like to thank Donald J. Zack and Arumugam Nagalingam (Glaucoma Center for Excellence, Wilmer Eye Institute, Johns Hopkins University School of Medicine) for providing the hiPSC-derived RGCs used in this work and William Yutzy (Glaucoma Center for Excellence, Wilmer Eye Institute, Johns Hopkins University School of Medicine) for helping us with the image analysis.

## Conflict of interest disclosure – data availability

The authors declare no competing financial interest. Furthermore, the authors confirm that the data supporting the findings of this study are available within the article [and/or] its supplementary materials.

## Ethics approval statement

Postmortem human eyes were obtained from Biobank Antwerpen, (Antwerp, Belgium; ID: BE 71030031000) and Ghent University Hospital. Protocols were approved by the Ethical Committee of Ghent University Hospital (dossier number: B670201837281; EC/2018/1071).

## Toc Text

Glaucoma, the leading cause of irreversible blindness, is linked to retinal ganglion cell (RGC) loss. Our research refines photodisruption, a minimally invasive technique that locally ablates the inner limiting membrane to improve stem cell-derived RGC transplantation. This technique improves donor cell migration and integration, paving the way for potential vision restoration through advanced cell replacement therapy.

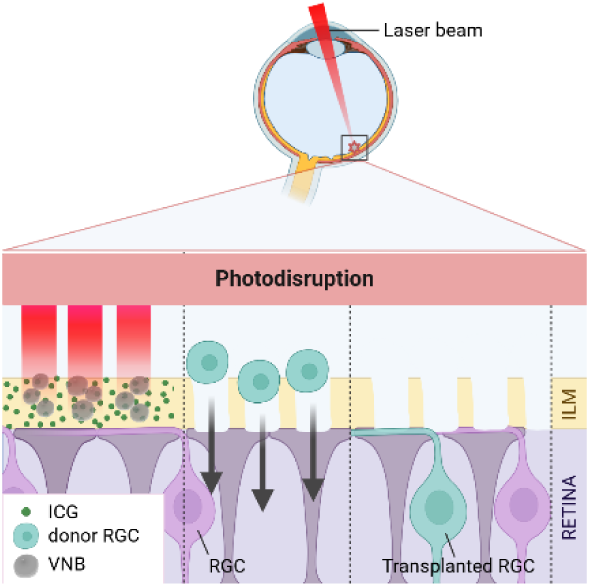

