## Supplementary figures for "Photodisruption of the inner limiting membrane promotes retinal engraftment of stem-cell derived retinal ganglion cells"

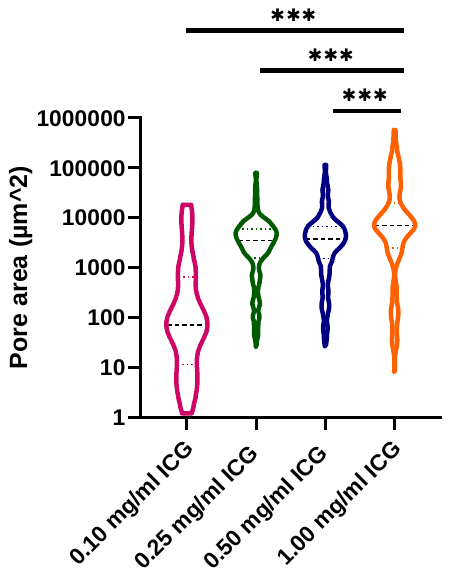

**Figure 1:** Quantification of the pore area for varying ICG concentrations. Increasing ICG concentrations resulted in elevated pore diameter and pore area. *p ≤ 0.05, **p ≤ 0.01, ***p ≤ 0.001, NS: p > 0.05.

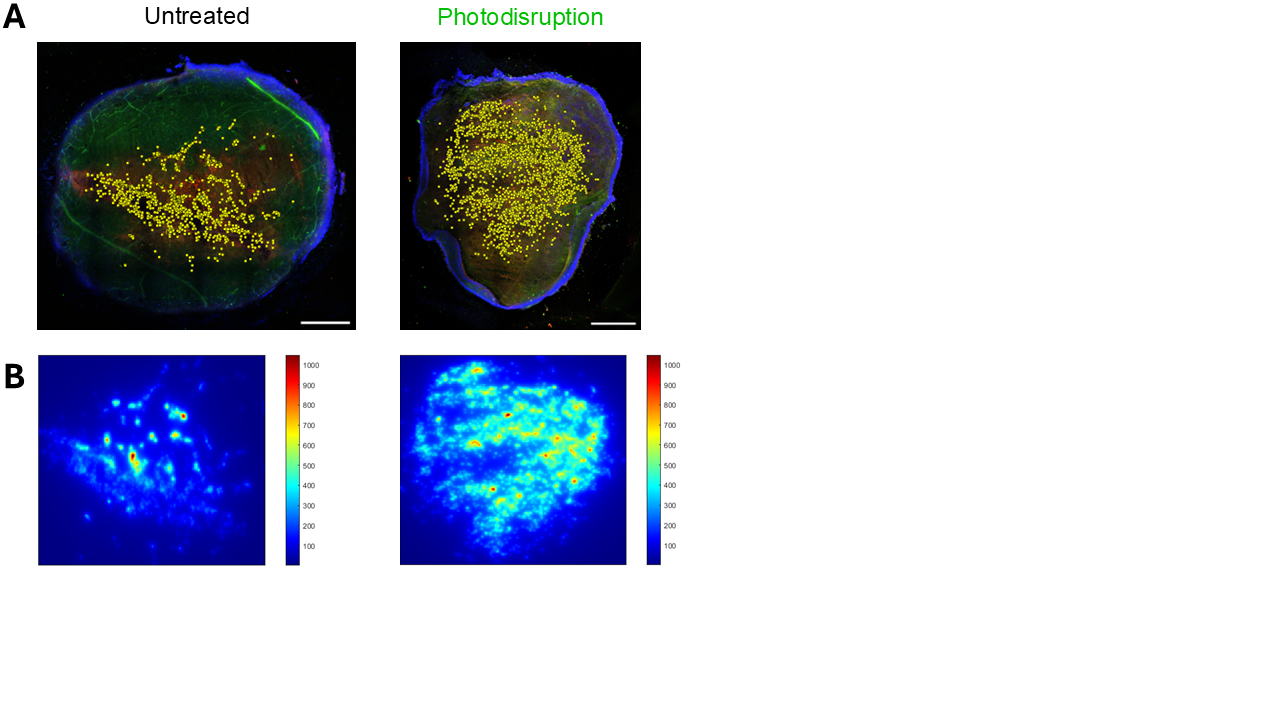

**Figure 2:** Representative confocal images of untreated and ILM disrupted bovine flatmounts, 7 days after transplantation with 20.000 hiPSC-derived RGCs expressing TdTomato (red). Flatmounts were stained with Human Nuclei (green) to identify the hiPSC-derived RGCs and Hoechst (blue) to visualize all nuclei (A). From the coordinates of the donor RGCs, density heat maps were generated using MATLAB (B). Scale bars: 1 mm (A).

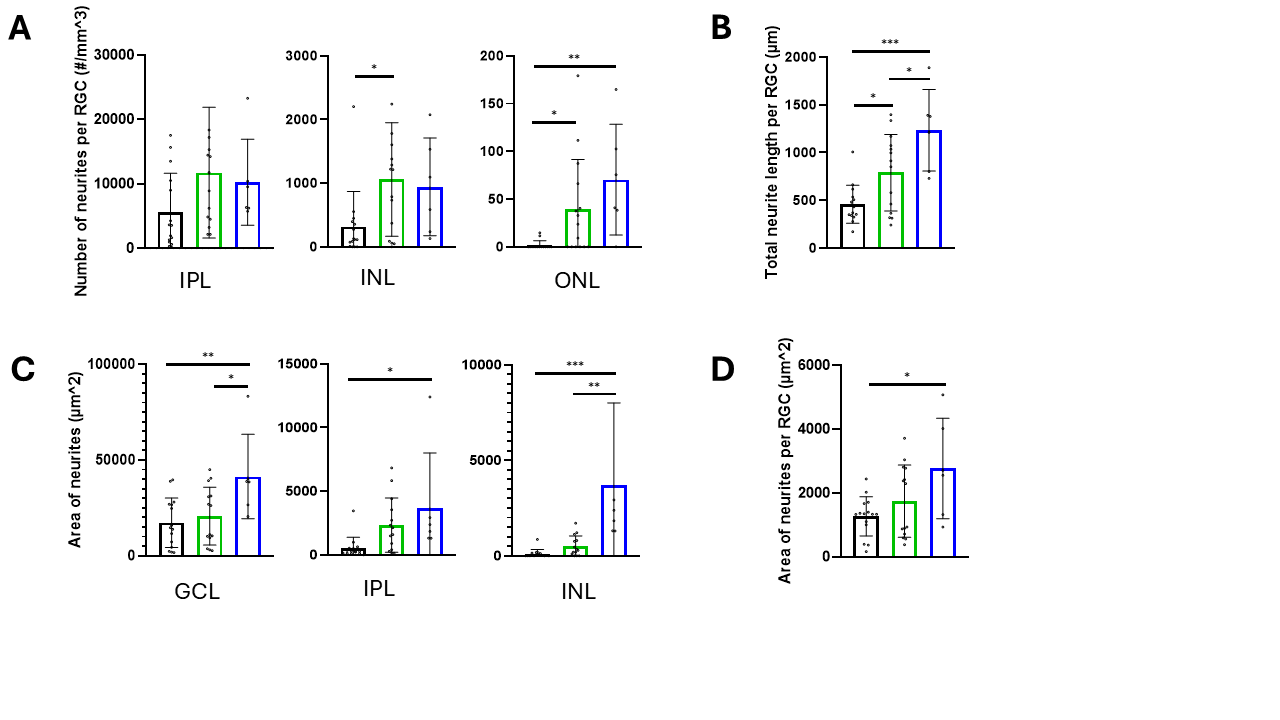

**Figure 3:** Quantification of the number of neurites per RGC in each cell layer. Using Fiji, the neurite localization was quantified by counting the number of neurites per RGC reaching each cell layer, the inner plexiform layer (IPL), the inner nuclear layer (INL) and the outer nuclear layer (ONL). Both treatments lead to more neurites per RGC in the IPL, along with the INL and ONL (A).The dendritic arbor area and the length of the neurites were semi-automatically determined in Imaris. Both the total neurite length per RGC (B) and the dendritic arbor area per RGC (C) increased following each ILM-disrupting treatment. *p ≤ 0.05, **p ≤ 0.01, ***p ≤ 0.001, NS: p > 0.05.

Untreated	Photodisruption	Collagenase

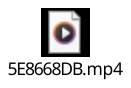

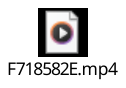

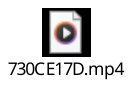

**Figure 4:** Representative movies of confocal Z-stack as visualized in 3D viewer in Fiji. One week after transplantation with hiPSC-derived RGCs expressing TdTomato (red), the flatmounts were stained with Hoechst (blue) to visualize all nuclei. The neurite localization was quantified by measuring the neurite depth and counting the number of neurites reaching each cell layer using 3D viewer or orthogonal view in Fiji.
